# Unconventional presentation of an immunodominant HLA-DR15-restricted nephritogenic proteinase 3 epitope implicated in vasculitis

**DOI:** 10.64898/2026.07.19.739468

**Authors:** C Lu, LR Shochet, JJ Lim, TJ Loh, S Goh, CM Jones, JB Zhang, L Allmannsberger, P Prakongtham, MT Tran, D Chisanga, J Ryan, L Fugger, M Moser, DE Jenne, HH Reid, NL La Gruta, AW Purcell, J Rossjohn, AR Kitching

**Affiliations:** Centre for Inflammatory Diseases, Monash University Department of Medicine, Monash Medical Centre, Clayton, Victoria 3168, Australia; Department of Nephrology, Monash Health, Clayton, Victoria 3168, Australia; Infection and Immunity Program, Biomedicine Discovery Institute and Department of Biochemistry and Molecular Biology, Monash University, Clayton, Victoria 3800, Australia; Department of Cancer Medicine, School of Translational Medicine, Monash University, Melbourne, Victoria 3004, Australia; Max-Planck-Institute of Biological Intelligence (formerly Max-Planck-Institute of Neurobiology), Am Klopferspitz 18, 82152 Planeg-Martinsried, Germany; Oxford Centre for Neuroinflammation, Nuffield Department of Clinical Neurosciences, and MRC Human Immunology Unit, Weatherall Institute of Molecular Medicine, John Radcliffe Hospital, University of Oxford, Oxford OX3 9DS, United Kingdom; Max PIanck Institute of Biochemistry, Department of Molecular Medicine, 82152 Martinsried, Germany; Department of Pediatric Nephrology, Monash Health, Clayton, Victoria 3168, Australia

## Abstract

Proteinase 3 (PR3) anti-neutrophil cytoplasmic antibody (ANCA)-associated vasculitis is associated with HLA-DR15, implicating HLA-DR15-restricted CD4⁺ T cell autoimmunity in the disease. However, the PR3 epitopes presented by HLA-DR15 and the autoreactive CD4⁺ T cell responses they elicit are undefined. Using humanized hPR3.DR15^+^ mice, we identified PR3_216-231_ as an immunodominant human PR3-derived CD4⁺ T cell epitope presented by HLA-DR15. Immunopeptidomic profiling confirmed the natural processing and HLA-DR15-restricted presentation of PR3_216-231_. Structural analyses of HLA-DR15-PR3_216-231_ complexes revealed an unconventional mode of HLA class II antigen presentation, in which a 10 amino acid peptide core occupies the 9 amino acid binding groove, causing a central kink in the bound peptide. PR3_216-231_-specific CD4⁺ T cells from both hPR3.DR15^+^ mice and patients with PR3-ANCA-associated vasculitis exhibited convergent T cell receptor features, and patients with active vasculitis displayed a clonally expanded, PR3_216-231_-specific CD4⁺ TCR repertoire. Immunization of hPR3.DR15 mice with PR3_216-231_ induced cell-mediated glomerulonephritis characterized by increased renal infiltration of CD4⁺ T cells and macrophages together with segmental glomerular necrosis. These findings identify a nephritogenic PR3-derived CD4⁺ T cell epitope presented by HLA-DR15 in ANCA-associated vasculitis and define a new extended core binding pattern, broadening our understanding of HLA class II peptide presentation.

## Introduction

Anti-neutrophil cytoplasmic antibody (ANCA)-associated vasculitis (AAV) is a life-threatening systemic autoimmune disease characterized by necrotizing inflammation of small blood vessels, with an incidence of 5-30 per million population per year (*1*). The kidneys and lower respiratory tract are common and critical targets, with renal and respiratory involvement occurring in approximately 65% and over 50% of cases, respectively. (*1, 2*). Untreated, AAV carries a high mortality and even with treatment, mortality is increased 2.7-fold compared with the general population (*3*). Unlike many autoimmune diseases, there is no female predominance. There are two major antigenic specificities, myeloperoxidase (MPO) and proteinase 3 (PR3), with PR3-AAV representing a distinct entity characterised by granulomatous inflammation and high relapse rates (*4*). Current immunosuppressive treatments have improved clinical outcomes, but do not completely prevent relapse and have significant side effects (*5*). Furthermore, there are no reliable biomarkers that define disease remission and predict relapse, leading to both unnecessary continuation of treatment and premature withdrawal of therapy, with consequent risks of over- and under-treatment (*6*).

While ANCAs can activate neutrophils, triggering neutrophil recruitment to vulnerable vascular beds, respiratory burst and degranulation, and endothelial damage (*7–9*), autoreactive CD4⁺ T cells also play a critical role in the pathogenesis of AAV (*10–12*). HLA class II alleles are important heritable risk factors for PR3-AAV, highlighting antigen presentation and autoreactive CD4⁺ T cells as mechanistic determinants of disease susceptibility (reviewed in ref (*13*)). Specifically, HLA-DPB1*04:01 (HLA-DP4) confers risk, with reported odds ratios (OR) of 3.4-6.4 across different Caucasian cohorts (*13–15*), and PR3-reactive CD4⁺ T cells have been detected and characterized in peripheral blood of HLA-DP4⁺ PR3-AAV patients (*10*). Similarly, HLA-DRB1*15:01 (HLA-DR15) represents another genetic risk allele, with an OR of 2 in Caucasian cohorts rising to 73 in African Americans (*15, 16*). However, the specific PR3 epitope(s) presented by HLA-DR15, the basis of peptide presentation, the clonality of DR15- restricted PR3-reactive CD4⁺ T cells, and the nephritogenicity of the epitope(s) are unknown.

Deciphering the mechanisms underpinning T-cell autoimmunity requires identifying the specific driving autoantigens and their interaction with risk-conferring HLA molecules. AAV stands out as a powerful and highly tractable clinical model to resolve these knowledge gaps because its autoantigens are well-defined, with almost all patients exhibiting autoreactivity to only one of PR3 or MPO. However, the sequence similarity between human and murine PR3 is only 69% (*17*), meaning that *in vivo* systems have not been able to model human anti-PR3 autoimmunity. Here we addressed this limitation by generating a new humanized transgenic mouse strain (hPR3.DR15⁺) that is knocked in for human PR3, deficient in murine MHC Class II, and transgenic for HLA-DRA1*01:01 and HLA-DRB1*15:01.

Using these humanized mice and samples from patients with PR3-AAV, we identify a single immunodominant HLA-DR15-related CD4⁺ T cell epitope (PR3_216-231_) that is naturally processed and presented by APCs from hPR3.DR15^+^ mice. Structural studies reveal an unconventional interaction with HLA-DR15, where the nine amino acid groove within HLA-DR15 accommodates a 10-residue core (PR3_219-228_). Single-cell paired TCRαβ TCR sequencing reveals similarities between that peptide-specific CD4⁺ TCRs in mice and HLA-DR15⁺ patients, and that patients with active vasculitis display a more restricted TCR repertoire than those in remission. We establish a role for PR3_216-231_ *in vivo*, with immunization resulting in nephritogenic autoimmunity and nephritis. Our studies establish a definitive, antigen-specific framework for understanding the pathogenesis of PR3-AAV, offering a foundation for the development of biomarkers that include both T cell and B cell anti-PR3 autoimmunity, as well as for epitope- and T cell-targeted immunotherapies.

## Results

### Human PR3_216-231_ is the only HLA-DR15-restricted PR3-derived CD4^+^ T cell epitope

Mice knocked in for human PR3, expressing exon 1 of mouse *Prtn3* (mPR3_1-26_) and exons 2-5 of human *PRTN3* (hPR3_21-256_) were used for these studies (hPR3^+/+^ mice). The mouse exon 1 derived sequence with the major portion of intron 1 was used to ensure fidelity of PR3 expression and processing (fig. S1). As exon 1 transcribes only the murine signal peptide, the protein represents the complete human proform of the hPR3 gene. Analyses confirmed the expression of hPR3 and its catalytic function in hPR3^+/+^ mice (fig. S2). Human PR3 mice were crossed with mice deficient in all elements of murine MHC-II and transgenic for HLA-DRA1*01:01 and HLA- DRB1*15:01 (*18*), generating mice expressing both human PR3 and the HLA-DR15 heterodimer (hPR3.DR15^+^ mice).

We defined the critical CD4^+^ T cell PR3 epitopes using a previously validated strategy (*11, 18*). We first immunized hPR3.DR15^+^ mice with three pools of hPR3 16-mers from an overlapping human PR3 peptide library encompassing the amino acid sequence inferred from *PRTN3* exons 2-5 (hPR3_21-256_, table S1), with each pool being fifteen 16- mer peptides overlapping by 11 amino acids. On restimulating lymphocytes from draining lymph nodes (dLNs) 10 days after immunization, substantial autoreactivity to human PR3 peptides was found exclusively within the carboxy terminal third of hPR3, pool 3 (hPR3_171-256,_ Fig. 1A). Separate experiments immunizing mice with peptides derived from either mouse or human *Prtn3/PRTN3* exon 1 (table S2) did not result in substantial autoreactivity on lymphocyte restimulation (fig. S3). While T and B cell autoreactivity to “mirror image” complementary PR3 sequences has been reported (*19, 20*), immunizing hPR3.DR15^+^ mice with complementary human PR3 peptide pools (table S3 and fig. S4A) did not result in consistent T cell reactivity to these peptides (fig. S4, B to D). Thus, the only substantial CD4^+^ T cell reactivity to hPR3 is found in the PR3_171-256_ region.

**Fig. 1.**
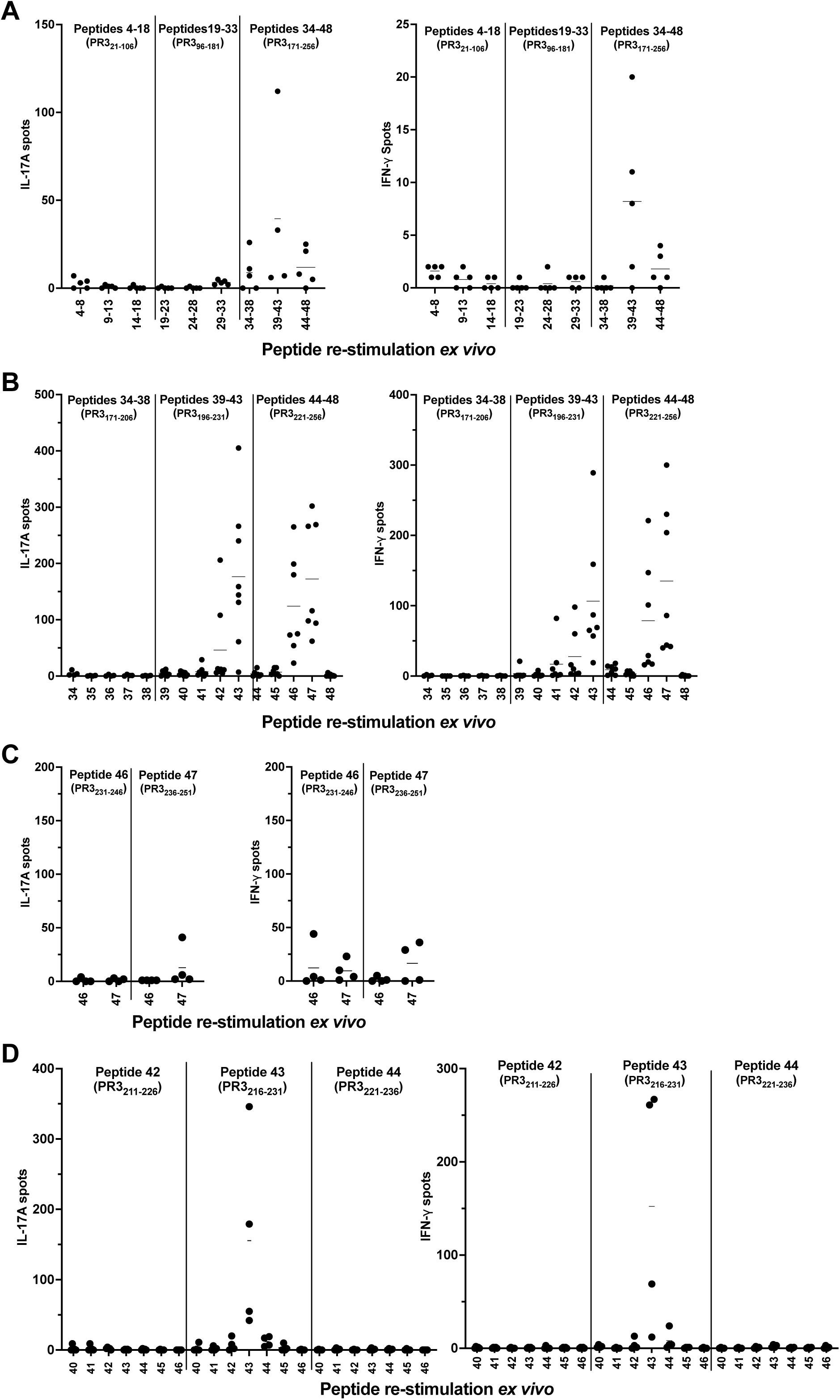
Defining PR3_216-231_ as the key PR3 T cell epitope in the context of HLA-DR15. hPR3.DR15^+^ mice were immunized subcutaneously at the base of tail on day 0 with hPR3 peptide pools or individual hPR3 peptides emulsified in Freund’s complete adjuvant (FCA). On day 10, mice were humanely euthanized and draining (inguinal and para-aortic) lymph nodes were harvested and restimulated *ex vivo* with the corresponding peptide pools or individual peptides. Antigen-specific T cell responses were quantified by IL-17A and IFN-γ ELISPOT assays. (**A to D**) IL-17A and IFN-γ ELISPOT analyses following: (**A**) Immunization with three hPR3 peptide pools (peptides 4-18, 19-33 and 34-48) and restimulation with the corresponding sub-pools; (**B**) Immunization with three sub-pools (peptides 34-38, 39-43 and 44-48) and restimulation with the corresponding individual peptides; (**C**) Immunization and restimulation with peptides 46 or 47; and (**D**) Immunization with peptide 42 or 43 followed by restimulation with peptides 40-46 individually. Horizontal bars indicate means for each group (n = 4 to 7 per group).

To pinpoint the T cell reactivity within PR3_171-256_ (pool 3), we immunized mice with one of 3 sub-pools from pool 3: peptides 34-38 (hPR3_171-206_), peptides 39-43 (hPR3_196-231_), and peptides 44-48 (hPR3_221-256_), and re-stimulated lymphocytes with each individual peptide from this pool. Peptides spanning PR3_211-231_ (peptides 42 and 43) and PR3_231-251_ (peptides 46 and 47,) were autoreactive (Fig. 1B). As peptide 43 (hPR3_216-231_, IDSFVIWGCATRLFPD) is the last peptide from the peptide 39-43 sub-pool, we immunized mice with a further sub-pool (peptides 41-45, hPR3_206-241_) to ensure that peptide 43 (hPR3_216-231_) was flanked on either side by any relevant amino acids (fig. S5A). Restimulation by peptide 43 (hPR3_216-231_) induced the strongest response. We then immunized mice with single peptides, with peptide 46 and peptide 47 inducing modest and inconsistent responses (Fig. 1C). Mice immunized with peptide 43 (hPR3_216-231_) responded to peptide 43 itself (IDSFVIWGCATRLFPD, Fig. 1D). T cells from mice immunized with recombinant hPR3 responded to peptide 43, but not to peptide 47 or a negative control peptide (peptide 17, fig. S5B). As these studies systemically define hPR3_216-231_ (peptide 43 IDSFVIWGCATRLFPD) as the only T cell epitope inducing strong and consistent autoimmunity when presented by HLA-DR15, further studies focused on this peptide (hPR3_216-231_).

As some T cell epitopes can induce B cell responses, hPR3.DR15^+^ mice were repeatedly immunized with hPR3_216-231_ and B cell responses assessed after 42 days. B cell ELISPOT showed reactivity against hPR3_216-231_, and hPR3_216-231_ antibodies were found in all mice (fig. S6A). Although low titre anti-hPR3 antibodies were present in most mice, B cell ELISPOTs to hPR3 were negative (fig. S6B) and these antibodies did not induce the characteristic cANCA pattern in human neutrophils (fig. S7). Therefore, while hPR3_216-231_ does induce anti-PR3 humoral responses, it is not the major PR3 B cell epitope.

Typically, 12-mers induce CD4^+^ T cell recall responses, although recall responses are stronger using 16 to 20-mer peptides. However, after immunizing mice with PR3_216-231_, none of the five 12-mers within the 16-mer PR3_216-231_ epitope induced recall responses to hPR3_216-231_ (Fig. 2A). This finding was supported by fluorescence polarisation-based competition assays, where these 12-mers bound only weakly to HLA-DR15 (Fig. 2B). Responses in 16-mer PR3_216-231_ immunized mice to truncated 13-15-mers were reduced, particularly 13-mers lacking the 3 amino acids at either the amino or carboxy terminals of PR3_216-231_ (Fig. 2C).

**Fig. 2.**
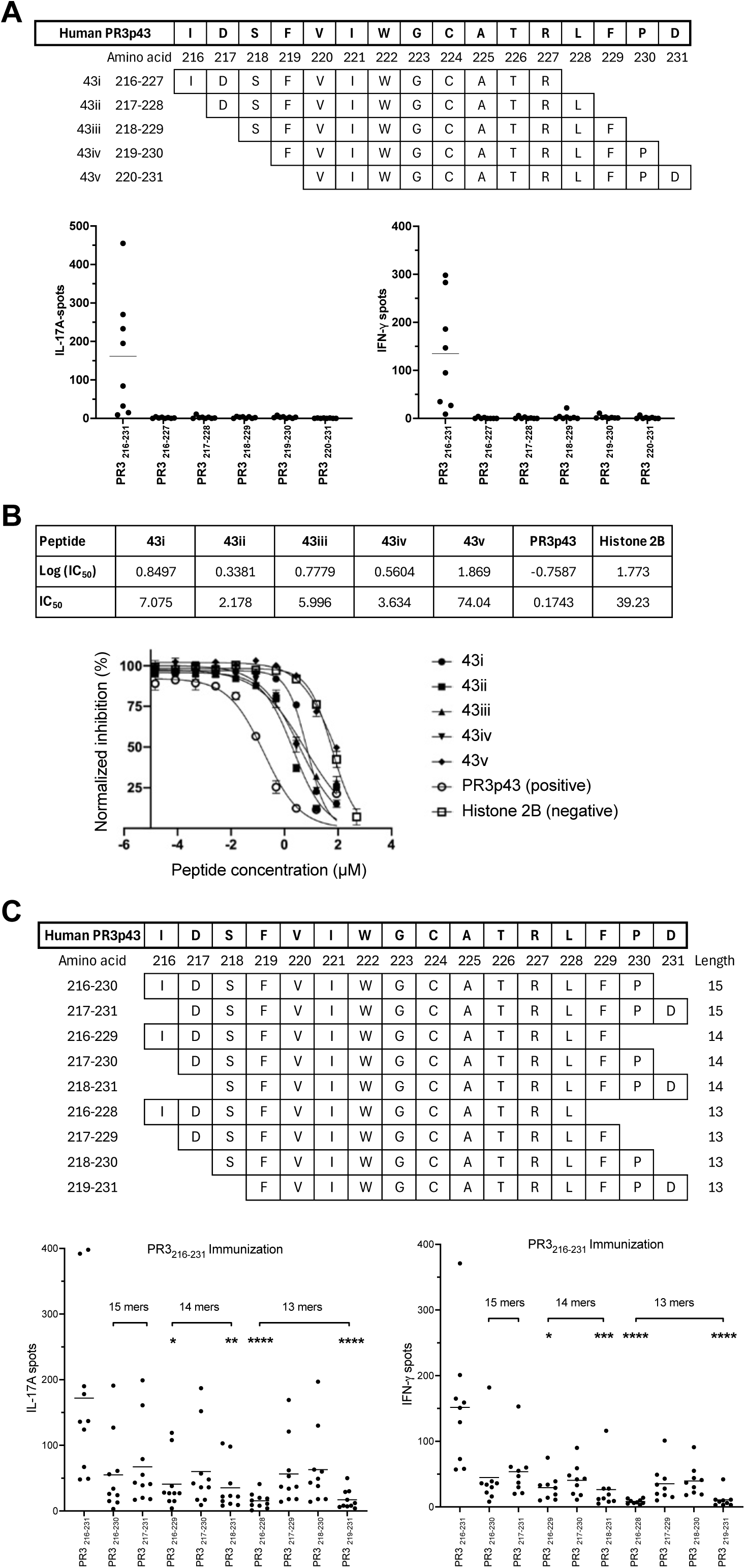
Production of IFN-γ and IL-17A is reduced or abolished in response to 12-mer or 13-15-mer derivatives of PR3_216-231_. (**A**) Comparison of antigen-specific T cell responses to hPR3 peptide 43 (PR3_216-231_) and its 12-mer derivatives (43i/PR3_216-227_, 43ii/PR3_217-228_, 43iii/PR3_218-229_, 43iv/PR3_219-230_ and 43v/PR3_220-231_) by IL-17A and IFN-γ ELISPOT assays performed 10 days post peptide immunization. (**B**) HLA-DR15 binding capacity of 12-mer hPR3 peptide 43 derivatives assessed by fluorescence polarisation-based competition assay. hPR3 peptide 43 and Histone 2B were the positive and negative control peptides, respectively. (**C**) Comparison of antigen-specific T cell responses to hPR3 peptide 43 (PR3_216-231_) and its 13-15-mer derivatives (15-mer: PR3_216-230_, PR3_217-231_; 14-mer: PR3_216-229_, PR3_217-230_, PR3_218-231_; 13-mer: PR3_216-228_, PR3_217-229_, PR3_218-231_, PR3_219-231_) by IL-17A and IFN-γ ELISPOT assays performed 10 days post peptide immunization. Horizontal bars indicate means (A and C, n = 8 to 10 per group) or means ± SD (B) for each group. In Panel C, statistical analyses used Kruskal-Wallis test with Dunn’s multiple comparisons test, comparing each truncated peptide with PR3_216-231_. \**P* < 0.05, \*\**P* < 0.01, *** *P* < 0.001, \*\*\*\**P* < 0.0001.

### PR3_216-231_ is naturally processed and presented by HLA-DR15 expressing APC

We next examined hPR3-derived peptides presented by HLA-DR15 molecules on antigen-presenting cells from freshly isolated splenocytes and lymph node cells from naïve hPR3.DR15^+^ mice, incubated either alone (untreated control) or with heat-inactivated recombinant hPR3 at concentrations of 20 μg/mL or 100 μg/mL. As hPR3.DR15^+^ mice express only one HLA allomorph, peptide-HLA-DR15 complexes were affinity purified using a pan-HLA-DR antibody and HLA-DR15-bound peptides pre-fractionated by RP-HPLC prior to mass spectrometry analysis. Raw data were searched using PEAKS Online against a mouse protein sequence database appended with the human PR3 protein sequence obtained from UniProt.

We detected 6,689 peptides derived from 1,263 source proteins in the untreated control, 4,749 peptides from 971 source proteins in the 20 μg/mL group, and 5,640 peptides from 1,119 proteins in the 100 μg/mL PR3 treated groups (Fig. 3A). As expected for HLA class II ligands, bound peptides ranged from 10 to 20 amino acids in length, with those containing 14 or 15 residues being most commonly identified (Fig. 3B). HLA-binding motif analysis of the identified 13-17-mer peptides revealed similar 9-mer core motifs across the three groups and was consistent with the expected HLA- DR15 binding motifs (Fig. 3C).

**Fig. 3.**
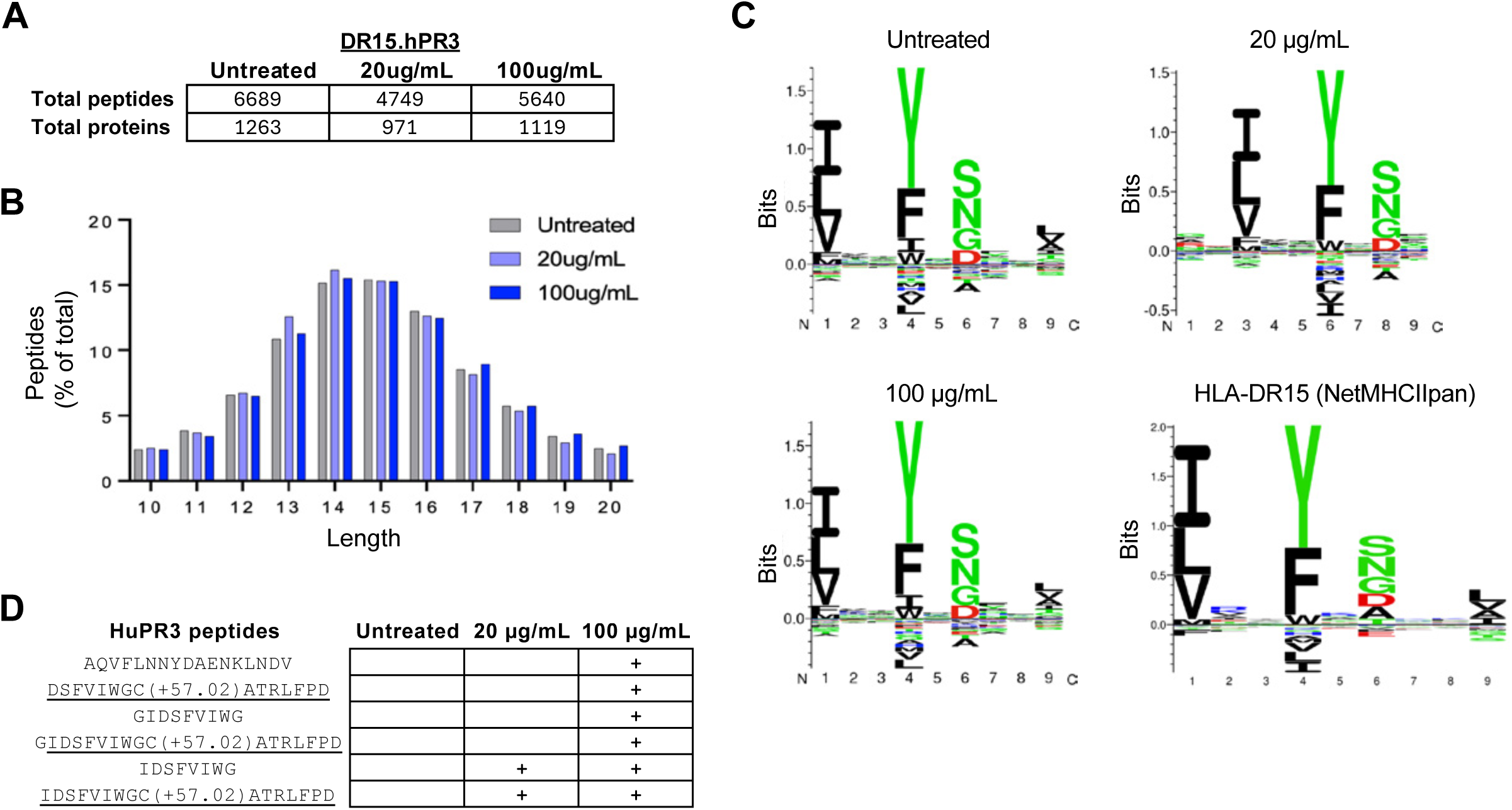
Immunopeptidome profile of HLA-DR-eluted peptides presented by antigen-presenting cells from hPR3.DR15^+^ mice. Freshly isolated splenocytes and lymph node cells from naïve hPR3.DR15^+^ mice were incubated either alone (untreated control) or with heat-inactivated recombinant hPR3 protein at concentrations of 20 μg/mL or 100 μg/mL prior to immunopeptidome analysis using mass spectrometry. (**A**) Total numbers of eluted peptides and corresponding source proteins identified. (**B**) Percentage distribution of peptides 10-20 amino acids in length. (**C**) GibbsCluster motif analysis of identified 13-17-mer peptides. (**D**) Identified hPR3-derived peptides.

Among the peptides detected, six were derived from hPR3 (Fig. 3D) originating from only two regions of the molecule (PR3_103-119_ and PR3_215-231_). No hPR3 peptides were detected in untreated control antigen presenting cells, two peptides were detected in cells incubated with both 20 μg/mL and 100 μg/mL hPR3, and four additional peptides were identified only after incubation with 100 μg/mL hPR3. Of these six peptides, five were derived from the PR3_215-231_ region, with the remaining peptide, PR3_103-119_ found only when cells were incubated with high concentrations of PR3. The PR3_216-231_ peptide, containing a carbamidomethylated cysteine deliberately introduced during sample preparation [IDSFVIWGC(+57.02) ATRLFPD], was detected in both the 20 µg/mL and 100 µg/mL treated cells. The PR3_103-119_ region is unlikely to be immunogenic, as immunization of DR15.hPR3^+^ mice with recombinant hPR3, peptide 43/PR3_216-231_ and peptides covering the PR3_103-119_ region (peptide 20/PR3_101-116_ and peptide 21/PR3_106-121_) elicited T cell responses only to PR3_216-231_ (table S1 and fig. S8).

Together, these findings demonstrate that antigen-presenting cells of hPR3.DR15^+^ mice process hPR3 and predominantly present the immunogenic PR3_216-231_ peptide.

### Non-canonical presentation of PR3_216-231_ by HLA-DR15

Given the immunodominance of PR3_216-231,_ the poor binding of 12-mers derived from PR3_216-231_ and the inability of these 12-mers to induce recall responses, we hypothesised that the HLA-DR15/PR3_216-231_ complex structure would be atypical. As the hPR3_216-231_ peptide exhibits poor aqueous solubility, two glutamate residues were added to the C-terminus, generating the peptide IDSFVIWGCATRLFPDEE (hereafter termed PR3_216-231_-Cys) to improve solubility). As the cysteine residue at PR3_224_ could promote peptide aggregation through disulfide bond formation, we generated an analogue in which cysteine was substituted with α-aminobutyric acid (Abu), a non-natural amino acid lacking the reactive thiol group that is less prone to oxidation and intermolecular cross-linking (*21–23*). This analogue is referred to as PR3_216-231_-Abu (IDSFVIWG(Abu)ATRLFPDEE).

We determined the crystal structures of HLA-DR15 in complex with PR3_216-231_-Cys and PR3_216-231_-Abu at 2.9 Å and 3.0 Å resolution, respectively. Both structures exhibited well-defined and unbiased electron density throughout the peptide bound within the HLA-DR15 groove, enabling unambiguous modelling of the peptide conformation (Fig. 4, A and B, table S4). Unexpectedly, both PR3_216-231_-Cys and PR3_216-231_-Abu adopted a non-canonical binding register within the HLA-DR15 antigen-binding groove (Fig. 4, B and C), In contrast to the conventional 9-mer core peptide register observed in HLA class II complexes, in which anchor residues occupy the P1, P4, P6 and P9 pockets(*24, 25*), both PR3_216-231_-Cys and PR3_216-231_-Abu bound as a 10-mer core epitope (P1-P10; IDS<u>FVIWGCATRL</u>FPD, core epitope underlined), engaging the HLA-DR15 groove through anchor residues at P1, P4, P8 and P10 (Fig. 4, C to F). Structural superposition of the two complexes revealed a root-mean-square deviation (r.m.s.d.) of 0.12 Å over HLA-DR15 Cα atoms, demonstrating that substitution of the P6 cysteine with Abu had no detectable impact on the overall mode of peptide presentation (Fig. S9).

**Fig. 4.**
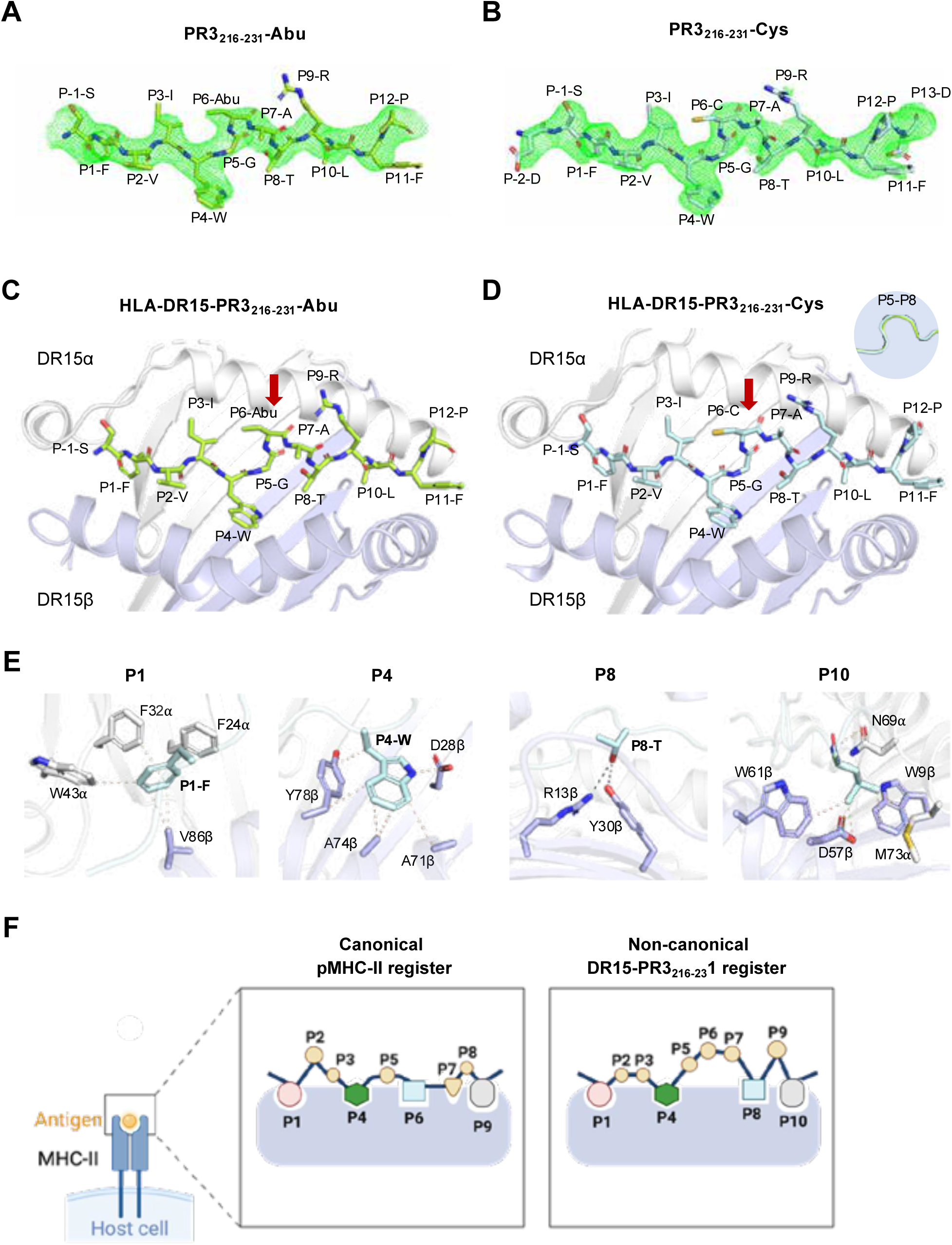
Non-canonical presentation of PR3_216-231_ by HLA-DR15. (A, B) Unbiased SA-omit σA-weighted Fo-Fc electron density maps (green mesh, contoured at 2.5σ) showing the bound peptides (A) PR3_216-231_Abu and (B) PR3_216-231_Cys within the HLA-DR15 antigen-binding groove. (C, D) Overall structures of HLA-DR15 presenting (C) PR3_216-231_Abu and (D) PR3_216-231_Cys. HLA-DR15 α− and β-chain are shown as gray and blue cartoons, respectively, while the peptides are depicted as sticks and coloured lemon (PR3_216-231_Abu) and cyan (PR3_216-231_Cys). Red arrows highlight the P5-P8 region, and inset views show the kinked peptide conformation that enables accommodation of the non-canonical 10-mer core epitope. (E) Detailed interactions between HLA-DR15 and PR3_216-231_Cys at the P1, P4, P8, and P10 anchor positions. Hydrogen bonds and van der Waals interactions are indicated by black and beige dashed lines, respectively. Amino acid residues are labelled using single-letter codes. (F) Schematic comparison of MHC class II peptide presentation modes, illustrating the conventional 9-mer core register (left) and the non-canonical HLA-DR15-PR3_216-231_ (right). Anchor residues are shown occupying the P1, P4, P6, and P9 pockets in the canonical register and the P1, P4, P8, and P10 pockets in the non-canonical register.

Notably, residues P5-P8 of both PR3_216-231_-Cys and PR3_216-231_-Abu peptides adopted a subtle kinked conformation that enabled the P4-Trp and P8-Thr side chains to engage their respective anchor pockets simultaneously (Fig. 4, C and D). To determine whether this unusual conformation was driven by the peptide sequence itself, we examined the interactions formed by the P1, P4, P8 and P10 anchor residues (Fig. 4E). P1-Phe and P4-Trp were deeply buried within extensive hydrophobic networks formed by both DR15 α-and β-chain, consistent with previously defined HLA- DR15 peptide binding-motifs (*26, 27*) (Fig. 4E). At the C-terminal end of the core epitope, P10-Leu was accommodated within a shallow hydrophobic pocket formed by Trp61β, Trp9β, Met73α, and Asp57α, corresponding to the canonical P9 pocket observed in other peptide MHC-II (pMHC-II) complexes (Fig. 4E). In contrast, P8-Thr occupied a previously uncharacterised shallow and polar pocket, where its hydroxyl group formed hydrogen-bonding interactions with Arg13β and Tyr30β (Fig. 4E). Together, the simultaneous anchoring of the bulky hydrophobic P4-Trp and the polar P8-Thr residues imposes a local distortion in the peptide backbone, giving rise to a P5-P8 backbone kink that underpins this non-canonical and unconventional 10-mer binding register.

This unconventional binding register defined in both HLA-DR15-PR3_216-231_ complexes alters the landscape available for TCR recognition. Specifically, residues at position P6, P7, and P9 protrude from the peptide-binding groove, creating an atypical solvent-exposed surface for TCR engagement (Fig. 4, C, D and F). In conventional HLA class II peptide presentation, residues at P2, P5, and P8 commonly represent the dominant TCR-contacting positions, as demonstrated in several TCR-pMHC-II structures associated with autoimmune disease including rheumatoid arthritis (*28–30*) and coeliac disease (*31, 32*) (Fig. 4F). In contrast, the shifted binding register adopted by hPR3_216-231_ repositions the exposed peptide side chains and remodels the antigenic landscape presented to T cells. Collectively, these findings reveal a previously unrecognised mode of peptide presentation by HLA-DR15 that is likely to generate a distinct TCR recognition surface and thereby influence autoreactive TCR selection and recognition.

### PR3_216-231-_specific mouse and human CD4 TCR repertories exhibit similar characteristics

We first characterized TCRαβ gene usage in PR3_216-231_-specific CD4⁺ T cells from PR3_216-231_-immunized hPR3.DR15^+^ mice. Consistent identification and isolation of PR3_216-231_-specific CD4⁺ T cells directly from dLNs and spleens of immunized mice was difficult. Therefore, epitope-specific T cells were enriched *ex vivo* by restimulating freshly isolated CD4⁺ T cells from dLNs and spleens of immunized mice with PR3_216-231_-Cys- or PR3_216-231_-Abu-pulsed bone marrow-derived dendritic cells (BMDCs) from hPR3.DR15^+^ mice. HLA-DR15-PR3_216-231_ tetramer staining confirmed clear tetramer⁺ populations after enrichment (Fig. 5A). Tetramer^+^ CD4⁺ T cells were single-cell sorted (Fig. 5B and fig. S10A), followed by paired TCRαβ sequencing. Detailed analysis of sequencing data was performed using the TCRdist algorithm (*33*). V-J gene pairing of each clonotype in the PR3_216-231_-specific hPR3.DR15^+^ mice CD4 TCR repertoire revealed preferred gene usage of TRAV6-7/DV9, TRBV19, TRBV20 and TRBV31 genes (fig. S11A) and clonal expansions characterized by TRAV6- 7/DV9_AJ42_TRBV31_BJ1-1, TRAV16_AJ53_TRBV19_BJ1-1, and TRAV13-2_AJ15_TRBV20_BJ2-5 gene combinations (Fig. 5C, fig. S11A and table S5).

**Fig. 5.**
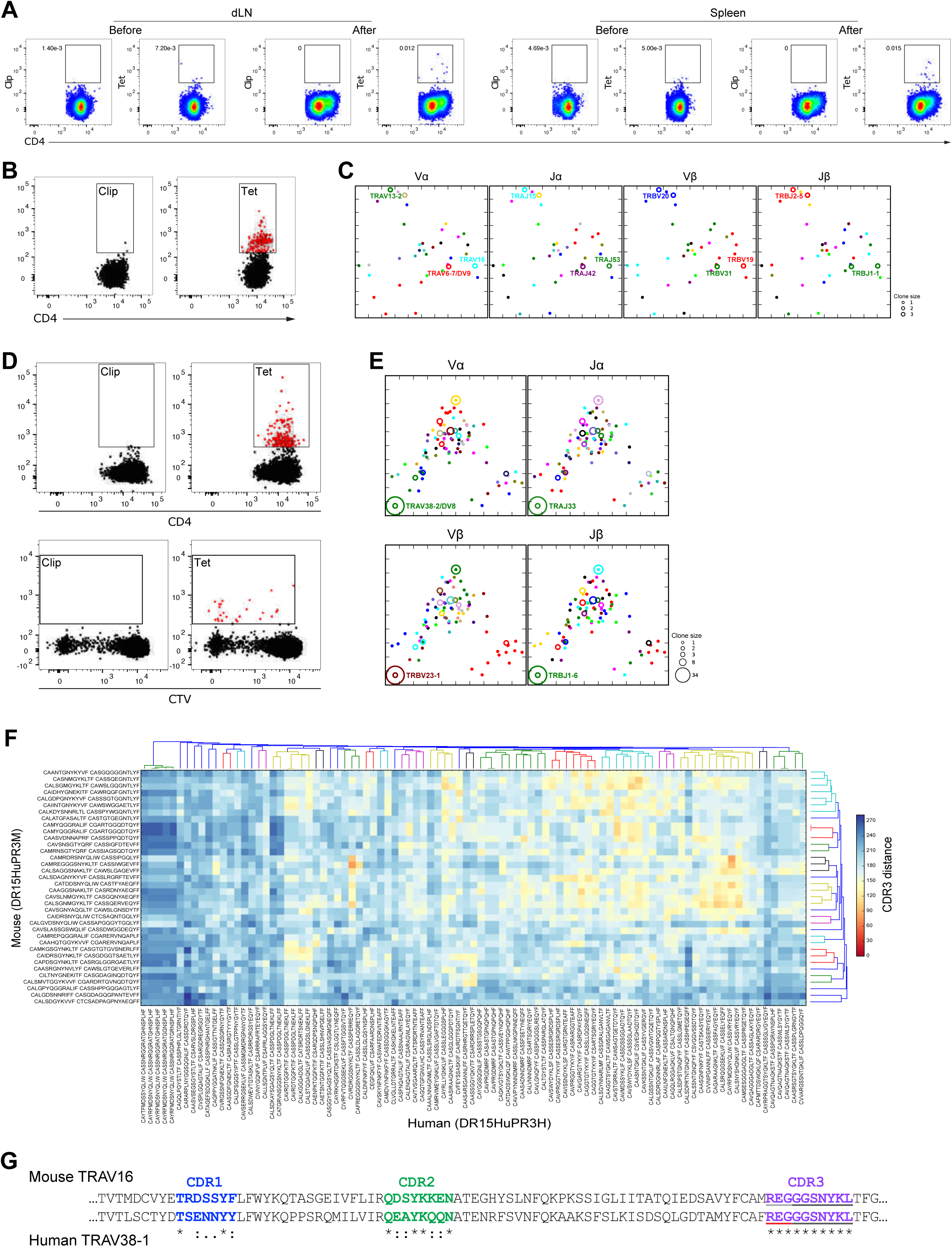
PR3_216-231_-specific CD4^+^ T cells exhibit conserved TCR features across hPR3.DR15^+^ mice and DR15^+^ PR3-AAV patients. (**A**) Flow cytometric analysis of DR15-PR3_216-231_-tetramer^+^ CD4^+^ T cells from dLNs and spleens of immunized hPR3.DR15^+^ mice before and after *in vitro* enrichment. (**B** and **D**) Single-cell sorting of tetramer^+^ cells from enriched PR3_216-231_-specific hPR3.DR15^+^ mouse CD4^+^ T cells (B), and from enriched PR3_216-231_-specific DR15^+^ PR3-AAV patient non-labelled (D, top panel) or CTV-labelled PBMCs (D, bottom panel). (**C** and **E**) Kernal principal component analysis (kPCA) projection of the TCRdist matrix within hPR3.DR15^+^ mice (C) and DR15^+^ PR3-AAV patients (E). TCR clones with clonal expansion are shown as circles with size reflecting clone size. Colours are based ononrepresent V or /J gene frequency within the repertoire as follows: from red (most frequent), green, blue, cyan, magenta followed by assorted colours for rare frequencies. (**F**) Heatmap of pairwise CDR3 distance between mouse and human TCR repertoires. CDR3 distance is calculated by removing CDR1 and CDR2 weighting from the original TCRdist method. Clonotypes are arranged via hierarchical clustering of TCRs within each repertoire, shown by dendrographs, to group similar clonotypes together. CDR3 alpha and beta sequences are annotated for each clonotype. (**G**) Clustal alignment of hPR3.DR15^+^ mouse TRAV16^+^ TCR and DR15^+^ PR3-AAV patient TRAV38-1^+^ TCR.TCRs. Highlighted CDR regions show sequence conservation: identical (*), highly conserved (:), semi conserved (.). Underlines indicate the source of CDR3 nucleotides, with light grey, dark grey and red denoting V, J and N regions, respectively.

Next, we defined the TCR repertoire of PR3_216-231_ specific CD4^+^ T cells in PR3-AAV patients expressing HLA-DRB1*15:01. A similar epitope-specific cell enrichment strategy was used for peripheral blood mononuclear cells (PBMCs) isolated from five HLA-DR15^+^ PR3-AAV^+^ patients (table S6), including three patients with active disease (A1, A2 and A3) and two patients in remission (R1 and R2). Paired TCRαβs from single-cell sorted HLA-DR15-PR3_216-231_ tetramer^+^ CD4^+^ T cells were sequenced (Fig. 5D, fig. S10, B and C). We retrieved 143 TCRs, representing 86 clones (table S7 and fig. S11B). Compared with mouse TCRs, clonal expansion was more pronounced among human TCRs, with the most dominantly expanded clone (clone frequency approximately 25%) using TRAV38-2/DV8_AJ33_TRB23-1_BJ1-6 gene combinations (Fig. 5E, fig. S11B and table S7). Detailed bioinformatic analysis of the TCR distance matrix, calculated using only CDR3 sequences to minimize gene-specific bias in TCRdist scores caused by the germline differences in mouse and human CDR1 and CDR2, revealed similar TCRs in the mouse and human repertoires (Fig. 5F). In particular, a human TRAV38-1^+^ clone not only shares an identical CDR3 region but also shows similarities in the germline-encoded CDR regions with the expanded mouse TRAV16^+^ clone (Fig. 5G), whose antigen specificity was validated by tetramer staining after in vitro TCR expression (fig. S12). Notably, while the “REG” motif in CDR3 region was V-gene-encoded by TRAV16 in the mice, in the human TRAV38-1^+^ clone it was encoded by N nucleotide additions (Fig. 5G), suggesting its importance in pHLA recognition. Collectively, these data show that antigen-driven selection shapes humanized mouse and human CD4⁺ TCRs specific for HLA-DR15-PR3_216-231_ in a similar manner, with the conserved CDR3 regions underscoring the relevance of these findings to human PR3-AAV.

### The HLA-DR15-PR3_216-231_ specific CD4 TCR repertoire is more restricted in active vasculitis

We next compared the composition of HLA-DR15-PR3_216-231_ specific CD4 TCR repertoires in HLA-DR15^+^ PR3-AAV^+^ patients with active vasculitis (n=3) and in remission (n=2). Analyses of clonotypes and clone size in both repertoires revealed that the expanded clonotypes were predominantly from patients with active vasculitis, with larger clonotypes dominating the active disease repertoire (Fig. 6A). Furthermore, the two most expanded clonotypes (TRAV38-2/DV8_AJ33_TRBV23-1_BJ1-6) and (TRAV20_AJ27_TRBV19_BJ1-2) were public, appearing in both remission and active disease patient repertoires (Fig. 6, A and B, table S7). Notably, in patients with active PR3-AAV, although the most expanded clones used public TCRs, private TCRs also contributed to clonal expansion (Fig. 6, A and B, fig. S13). Kernel principal component analysis (kPCA) of CDR sequence distances showed that HLA-DR15-PR3_216-231_ specific CD4^+^ TCR clonotypes in the active group displayed clustered signatures, indicative of closely related TCRs, whereas clonotypes from the remission group were more diverse and spatially dispersed (Fig. 6B). This was further reflected by the decreased nearest-neighbor (NN) distance and lower diversity within the active repertoire (Fig. 6C). Overall, defining and dissecting the HLA-DR15-PR3_216-231_ specific CD4 TCR repertoire reveals an expansion of closely related TCRαβ clonotypes during active disease, leading to a more restricted repertoire.

**Fig. 6.**
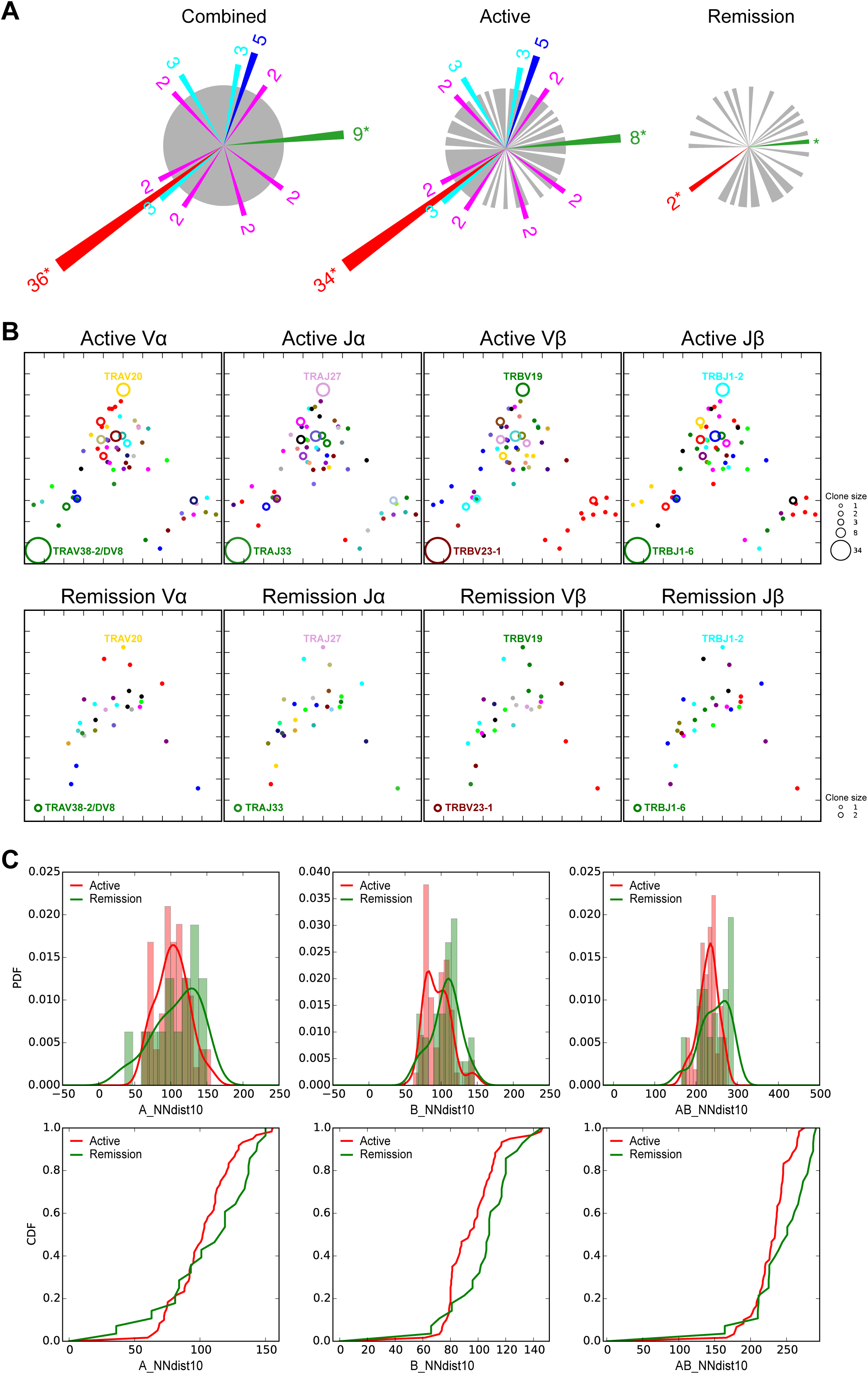
PR3_216-231_-specific CD4 TCR repertoire is restricted in DR15^+^ PR3-AAV patients with active disease. Paired TCRαβ repertoires of PR3_216-231_-specific CD4^+^ T cells isolated from DR15^+^ PR3-AAV patients with active disease and in remission. (**A**) Coxcomb plots show expanded (n > 1) and unique (n = 1) clones from DR15^+^ PR3-AAV patients with active disease and in remission. Clone size is color-coded, with red indicating the most expanded clone, followed by green, blue, cyan, magenta, and unique clones shown in grey. Asterisks (*) denote public clones. (**B**) Kernal principal component analysis (kPCA) projection of the TCRdist matrix within DR15^+^ PR3-AAV patients with active disease (top row) and remission (bottom row). TCR clones with clonal expansion are shown as circles with size corresponding to clone size. (**C**) The nearest neighbour distance (threshold 10^th^ percentile) distribution plot of the TCR repertoire from patients with active disease or in remission. Top row: Probability Density Functions (PDF) shown in bar and smoothed histograms. Bottom row: Cumulative Distribution Functions (CDF). Left to right: alpha chain, beta chain, combined).

### PR3_216-231_ immunization leads to glomerulonephritis

To determine the nephritogenic potential of anti-PR3 T-cell responses elicited by PR3 216–231 immunization *in vivo*, we used a T cell–dependent, antibody-independent glomerulonephritis model conceptually similar to an established anti-MPO model (*11, 12*). but adapted to PR3 as the autoantigen (Fig. 7A). PR3_216-231_ immunized hPR3.DR15^+^ mice lose tolerance to PR3, but do not develop nephritis, as PR3 is not present in glomeruli. To allow anti-PR3 T cell autoimmunity to mediate glomerular injury, mice were injected with a low dose of sheep anti-mouse glomerular basement membrane (GBM) globulin to induce transient glomerular neutrophil accumulation and glomerular PR3 deposition. Control mice were immunized with an irrelevant peptide (OVA_323-339_) prior to anti-mouse GBM antibodies.

**Fig. 7.**
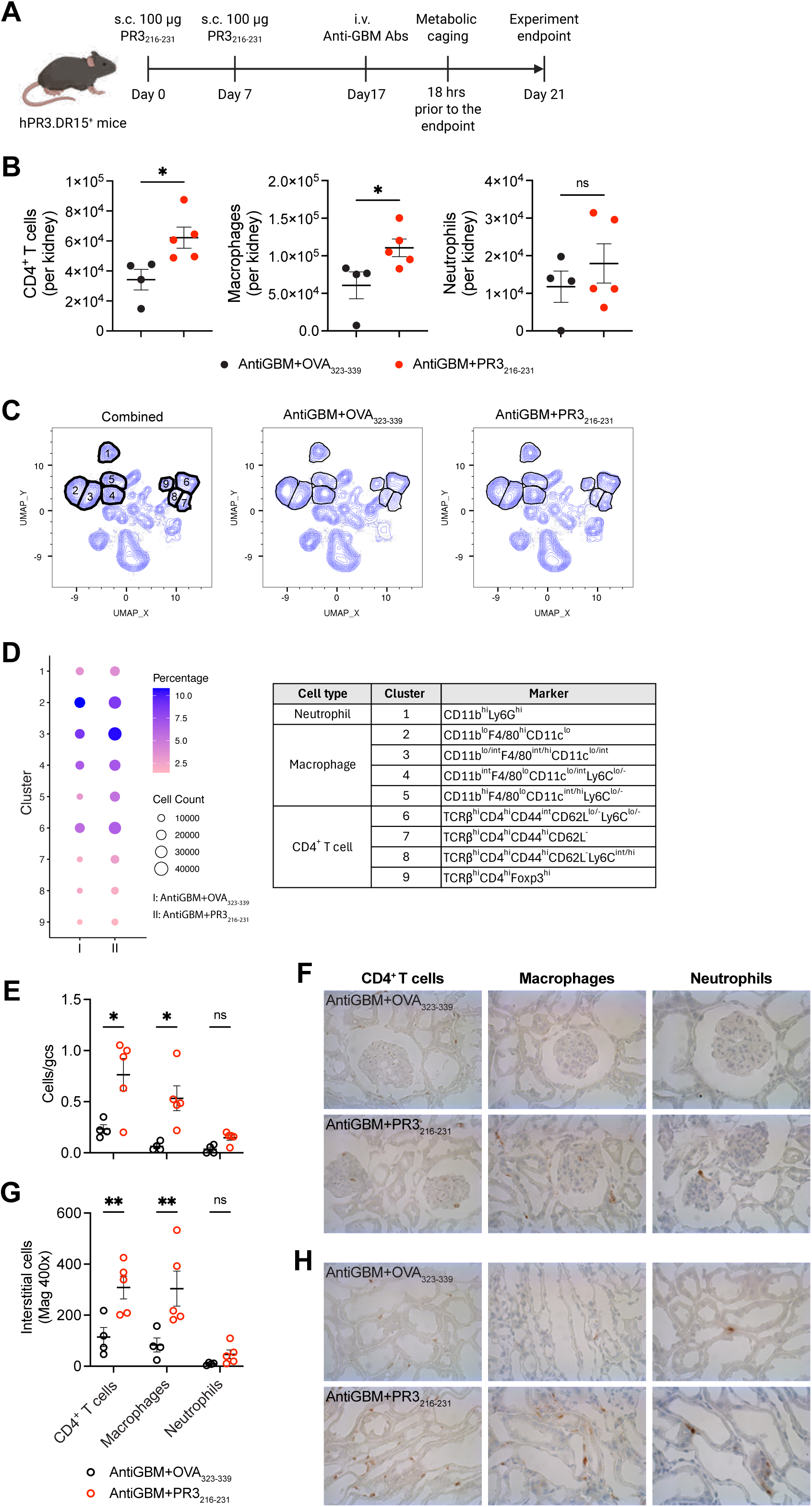
Immunization with PR3_216-231_ increases renal infiltration of CD4^+^ T cells and macrophages. DR15.hPR3 mice were immunized with PR3_216-231_ peptides (n=5) or the control peptide OVA_323-339_ (n=4) on days 0 and 7, followed by tail-vein injection of low-dose sheep anti-mouse GBM antibodies on day 17. Mice immunized with PR3_216-231_ peptides (AntiGBM + PR3_216-231_) and control mice (AntiGBM + OVA_323-339_) were euthanized on day 21. Kidney cells were isolated and analysed by high-parameter flow cytometry. (**A**) Experimental design for the assessment of the nephritogenicity of PR3_216-231_. (**B**) Flow cytometric quantification of CD4^+^ T cells, macrophages and neutrophils in the kidneys via flow cytometry. (**C**) UMAP analysis of renal CD45.2^+^ leukocytes, with clusters 1-9 indicated. (**D**) Dot plot summarizing the percentage and number of cells in each cluster, which was classified as neutrophils, macrophages or CD4^+^ T cells based on phenotypic markers. (**E** to **H**) Quantification and representative immunohistochemical staining of glomerular (**E** and **F**) and interstitial (**G** and **H**) CD4^+^ T cells, macrophages and neutrophils. Data are shown as mean ± SD. Statistical analysis was performed using Student’s *t* test (with Holm-Šídák’s correction for multiple comparisons). \**P* < 0.05; \*\**P* < 0.01. ns, not significant.

Flow cytometry analysis of intrarenal cellular infiltrates using unsupervised UMAP clustering four days after anti-mouse GBM globulin injection revealed, in PR3_216-231_ immunized mice, increased intrarenal infiltration of CD4^+^ T cells and macrophages, but not neutrophils, compared with control mice (Fig. 7B and fig. S14). Detailed analyses of UMAP-identified clusters of neutrophils, macrophages and CD4^+^ T cells also demonstrated that PR3_216-231_ immunization led to increased absolute numbers, with minimal changes in proportions, of macrophage subsets (clusters 2-5) and CD4^+^ T cell subsets (cluster 6-9), whereas there were only minimal differences in intrarenal neutrophils (cluster 1) (Fig. 7, C and D and Fig. S15). Immunohistochemical analyses supported these findings, with increased glomerular and interstitial infiltration of CD4^+^ T cells, and macrophages, but not neutrophils, in PR3_216-231_-immunized mice (Fig. 7, E to H). Moreover, PR3_216-231_-immunized mice developed glomerulonephritis (37.1 ± 14.3% of glomeruli affected) (Fig. 8A), characterized by segmental glomerular necrosis in some glomeruli (Fig. 8B), albeit without a significant increase in albuminuria (Fig. 8C). In contrast, glomeruli of control mice were largely unaffected, with only a small proportion (5.9 ± 2.1%) exhibiting mild lesions, reflecting the effects of anti-GBM globulin (Fig. 8D). Collectively, these findings demonstrate that CD4^+^ T cells induced by PR3_216-231_ immunization are nephritogenic and can mediate glomerular disease.

**Fig. 8.**
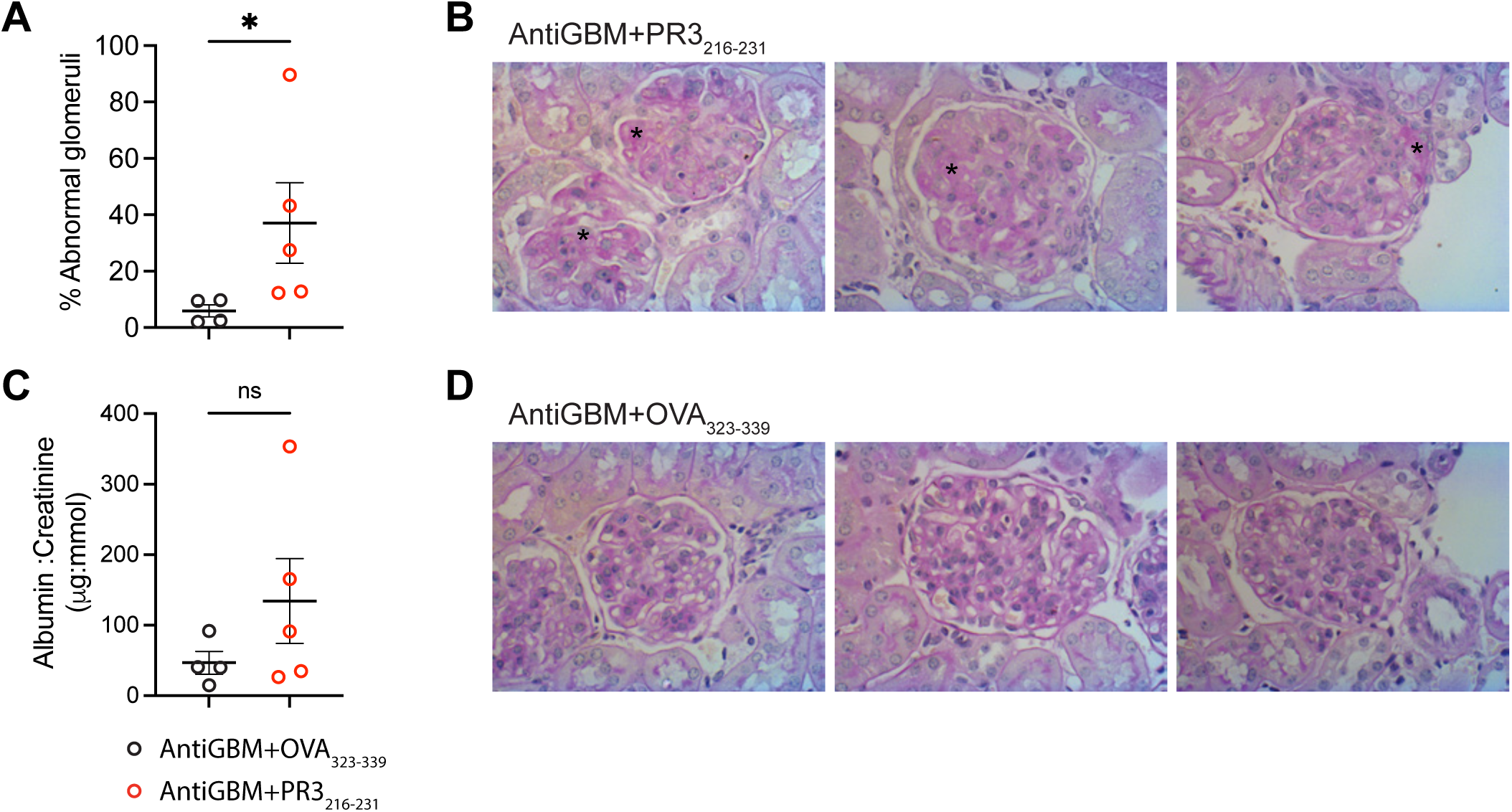
PR3_216-231_ immunization induces autoimmune nephritis. DR15.hPR3 mice were immunized with either OVA_323-339_ (control, n =4) or PR3_216-231_ (n = 5), and disease was triggered by recruiting neutrophils to glomeruli with low-dose sheep anti-mouse GBM antibodies. (**A**) Quantification of abnormal glomeruli on Periodic acid-Schiff (PAS)-stained, paraffin-embedded kidney sections. (**B**) Representative PAS-stained kidney sections from PR3_216-231_-immunized mice; Asterisks (*) indicate areas of glomerular necrosis. (**C**) Albumin to creatinine ratio as a measure of renal function. (**D**) Representative PAS-stained kidney sections from OVA_323-339_-immunized control mice showing largely unaffected glomeruli. Data are shown as mean ± SD. Statistical analysis was performed using Student’s *t* test. \**P* < 0.05. ns, not significant.

## Discussion

Loss of tolerance to the serine protease PR3 results in PR3-AAV, a type of autoimmune vasculitis that typically affects the kidneys and respiratory tract, but can involve any organ or tissue (*1*). Our studies define an immunodominant, nephritogenic PR3 CD4^+^ T cell epitope, with PR3_216-231_-specific CD4^+^ T cells found in both humanised mice and in humans with PR3-AAV. PR3_216-231_, binds to HLA-DR15 in an unconventional manner, with a 10-residue core accommodated across only 9 positions with within the HLA-DRA1*01:01/DRB1*15:01 binding cleft.

As HLA Class II differs substantially to murine MHC II, and as differences between murine PR3 and human PR3 have hindered past attempts to understand PR3-AAV (*17*), we generated mice expressing both human PR3 and HLA-DR15, one of the risk HLA allomorphs for PR3-AAV. To ensure fidelity of human PR3 expression, these mice express the signal peptide of mouse PR3 (mPR3_1-26_, homologous to hPR3_1-21_) that we determined was not immunogenic in the context of HLA-DR15 (fig S3). We used these hPR3.DR15^+^ mice to define a single immunodominant PR3 peptide (PR3_216-231_), which was presented by hPR3-pulsed DR15^+^ murine splenocytes. Immunization of hPR3.DR15^+^ mice with PR3_216-231_ induced proinflammatory PR3_216-231_ specific recall responses, and PR3_216-231_ stimulated hPR3 immunized murine lymphocytes. In DR15^+^ humans with PR3-AAV and active anti-PR3 autoimmunity (a positive serum PR3-ANCA at the time of sampling), we identified PR3_216-231_ specific CD4^+^ T cells using PR3_216-231_-DR15 tetramers, after *ex vivo* T cell culture with PR3_216-231._

HLA class II allomorphs are open ended, allowing binding of variable length peptides within a framework of 9 residues within the binding groove, with flanking peptides outside the HLA class II groove stabilising binding, modulating TCR affinity and affecting T cell activation (*34*). After defining PR3_216-231_ as the key 16-mer peptide, we sought, as in previous work, to shorten PR3_216-231_ to define the core binding sequence, typically consisting of 12 amino acids. However, 12 mers displayed a low affinity for HLA-DR15 and did not re-stimulate lymphocytes from immunized mice. Compared with restimulation using the full length PR3_216-231_, ELISPOT IL-17A and IFN-γ production was suboptimal using 13 to 15 mers. While the requirement for a longer peptide than anticipated for CD4^+^ T cell activation is unusual, it is not unprecedented (*35*), as amino acids within flanking region of at least one peptide has been shown to make contact with the TCR, meaning a longer peptide length is required for immunogenicity(*30, 36*).

Peptide antigen presentation by MHC class II molecule is a fundamental determinant for CD4^+^ T cell selection, repertoire formation and antigen recognition. The prevailing paradigm of MHC-II peptide presentation, established through decades of structural and biochemical studies, posits that peptides are accommodated within the MHC-II binding groove as a conserved 9-mer core epitope (*24, 25*). Regardless of peptide length, orientation, or flanking residues, peptide binding is typically stabilised through the engagement of anchor residues occupying the P1, P4, P6, and P9 pockets of the MHC-II groove (*24, 25*). Variations within this framework have been reported, including altered register, such as α3_135-145_ peptide from the non-collagenous domain of the α3 chain of type IV collagen critical to Goodpasture’s disease that binds to both HLA-DR1 and HLA-DR15 using distinct peptide registers (*37*). Similarly, post-translational modifications can influence peptide loading and HLA binding without fundamentally altering the canonical mode of presentation. For example, citrullination of self-peptides derived from fibrinogen, vimentin, and Tenascin C enhances binding to HLA-DR4 implicated in rheumatoid arthritis (*30, 38, 39*). Likewise, deamidation of gliadin peptides promotes binding to HLA-DQ2.5 in coeliac disease (*31*), whereas transpeptidation between proinsulin and C-peptide generates hybrid peptides that bind HLA-DQ8 in Type-I diabetes (*36, 40*). Despite these examples of altered peptide specificity and register usage, the resulting complexes retain the canonical 9-mer core architecture and engage the conserved P1, P4, P6, and P9 anchor pockets. Indeed, this mode of presentation has been observed across diverse HLA class II allomorphs and forms the foundation of current peptide-binding prediction algorithms and our understanding of CD4^+^ T cell immunity(*25*). More recently, an exception to the conventional binding orientation was described for cytomegalovirus-derived peptides presented by HLA-DP5, which bind in a reverse C-to-N orientation rather than the conventional N-to-C direction (*41, 42*). Nevertheless, these peptides still preserve the canonical 9-mer core architecture and utilise the same anchoring pockets within the MHC-II groove, underscoring the remarkable conservation of the structural principles governing MHC-II peptide presentation.

In this study, we identify a previously unrecognised mode of peptide presentation by HLA-DR15 that extends beyond established paradigms of MHC-II antigen presentation. Structural analysis revealed that the hPR3_216-231_ peptide adopts a subtle kink within its central backbone, enabling accommodation of a 10-mer core epitope within the HLA-DR15 binding groove. This non-canonical register is stabilised by anchor residues at P1, P4, P8, and P10, rather than the conventional P1, P4, P6, and P9 positions. The shifted register is driven by the simultaneous accommodation of a bulky P4-Trp residues and a polar P8-Thr within distinct binding pockets, resulting in a local distortion of the peptide backbone and repositioning of solvent-exposed residues available for TCR recognition. These findings reveal an unexpected degree of structural plasticity in HLA-DR15-mediated antigen presentation and suggest that HLA-DR15 can accommodate a broader range of peptide confirmations than previously appreciated. By generating distinct antigenic surfaces for TCR engagement, this alternative mode of peptide presentation may expand the repertoire of presented self-antigens and contribute to autoreactive T cell recognition in HLA-DR15- assoiacted autoimmune disease.

Using HLA-DR15-PR3_216-231_ tetramers we identified multiple PR3_216-231_ specific TCRs, initially from cells of PR3_216-231_ immunized hPR3.DR5^+^ mice and then from five humans with PR3-AAV and active autoimmunity. The requirement for *ex vivo* expansion of these peptide specific cells for clear and consistent tetramer-based identification is consistent with a previous HLA-DP4 based study in PR3-AAV (*43*). The similarities between some of the mouse and human TCRs validates our humanised mouse approach to PR3 epitope discovery and is consistent with structural requirements imposed on TCR selection and expansion by the HLA-DR15-PR3_216-231_ complex. Importantly, the two public clones detected and the closer relationships between TCRs in each of the three patients with active vasculitis compared with the two patients in remission (but with a positive serum PR3-ANCA) imply that a focussed PR3_216-231_ specific CD4^+^ T cell response may be important in disease activity. In addition, while identification of autoantigen specific T cells is challenging in clinical practice, our results imply that concurrent assessment of PR3-specific cellular and humoral immunity may have future value in assessing disease activity. Our findings are broadly consistent with a previous study of HLA-DP4 positive individuals with PR3-AAV that identified public PR3-reactive TCRs and conserved TCR features but did not define PR3 peptide specificity (*10*).

PR3-ANCA are important mediators of tissue injury in PR3-AAV. In addition to T cell reactivity, PR3_216-231_ immunization did induce anti-PR3 B cell responses. As conformational PR3 B cell epitopes in AAV may be more relevant than linear epitopes (*44*), PR3_216-231_ itself is unlikely to be a critical B cell PR3 epitope. However, immunising mice with PR3_216-231_ induced PR3_216-231_-specific autoantibodies and low titer PR3-ANCA, implicating this PR3 peptide in CD4^+^ T cell/B cell help in PR3-AAV. Exposure to PR3-derived complementary peptides has been implicated in humoral and cellular responses in PR3-AAV (*19, 20*) and linked to HLA-DR15 (*16*). However, immunization of hPR3.DR15^+^ mice with overlapping peptides derived from the complementary human PR3 sequence did not induce consistent primary T cell immune responses in immunized hPR3.DR15^+^ mice, findings that do not support the complementary PR3 hypothesis in loss of tolerance to PR3.

In AAV, T cells provide essential help for autoreactive B cells that themselves secrete ANCA. These autoantibodies activate primed neutrophils, causing them to actively lodge in vulnerable microvascular beds where they induce acute injury (*45*). They also release their autoantigen where it can be recognised by effector Th1 and Th17 antigen specific cells (*11, 46*) and humans with PR3-AAV exhibit expanded PR3-specific cytotoxic CD4^+^ T cells that participate in tissue injury (*10*). Immunising hPR3.DR15^+^ mice with PR3_216-231_ generated antigen specific effector CD4^+^ T cells that localised to kidneys after neutrophils had been transiently recruited by injection of low dose anti- basement membrane globulin. The resultant accumulation of CD4^+^ T cells and histological injury in PR3_216-231_ immunized mice provides proof of concept that PR3_216- 231_ induced CD4^+^ T cells are nephritogenic *in vivo*.

Genetic data implicate the abundance of PR3 in the risk of the development of autoimmunity to PR3 and PR3-AAV. In our studies no human PR3 peptides were found in resting hPR3.DR15^+^ splenocytes, with PR3_216-231_ and related peptides increasing when cells were cultured with human PR3, These observations are concordant with findings of two AAV GWAS identifying risk variants for PR3-AAV that increase the availability of PR3 to the immune system (*47, 48*). One variant, in *PRTN3* itself, enhances PR3 expression without altering its structure, while another in *SERPINA1* reduces levels of the PR3 inhibitor a1 anti-trypsin, compromising normal clearance and inactivation of PR3 and thus increasing the duration and intensity of autoantigen exposure. Both variants have been validated at the protein level (*48, 49*).

This is the second CD4^+^ PR3 epitope identified, with the other, PR3_225-239_ binding being selected from PR3 peptides that were predicted to bind strongly to HLA- DPA1*01:01/ DPB1*04:01 (HLA-DP4) (*43*). This sequence overlaps PR3_216-231_, suggesting that in PR3-AAV there is an epitope hot spot in this region of the autoantigen, as in MPO-AAV and Goodpasture’s disease, other forms of renal vasculitis (*37, 50*). Whether there is an additive or synergistic relationship between HLA-DR15 and HLA-DP4 in PR3-AAV in the risk or severity of disease is unclear. While autoimmunity is key in vasculitis activity and relapse, current tools measure only B cell autoimmunity to PR3. While PR3-ANCA has attracted attention both as a mediator of disease and as a biomarker, ANCA is not always concordant with disease activity and cannot be used to determine treatment options in an individual. Defining critical PR3 CD4^+^ T cell epitopes in PR3-AAV has implications for biomarker development, with the potential to combine humoral and cellular readouts of anti-PR3 autoimmunity to improve assessment of disease and prediction of relapse. Furthermore, our studies inform future antigen specific therapies for this autoimmune disease.

Our studies define the immunodominant and nephritogenic CD4^+^ T cell epitope, PR3_216-231_ in people with PR3-AAV who are HLA-DR15^+^. This peptide binds to HLA- DR15 in an unconventional, kinked manner, with a 10-residue peptide core accommodated across only nine positions of the HLA class II groove in an altered binding register, a feature that challenges current HLA epitope prediction algorithms that do not accommodate more than nine residues binding within the peptide groove. Furthermore, the delineation of this PR3 T cell epitope has implications both for our understanding of PR3-AAV pathogenesis and for the development of biomarkers and new therapies for vasculitis.

## Materials and Methods

### Mice

hPR3.DR15^+^ mice were generated by crossing humanized PR3 knock in mice (*51*) generated by Ozgene (Perth, Australia), with HLA-DR15 transgenic mice (expressing HLA-DRA1*01:01 and HLA-DRB1*15:01 and deficient in all murine MHC II genes) (*37, 52*). After crossing, hPR3.DR15^+^ were on the following background: 129S6/SvEv 50%, C57BL/10 25%, C57BL/6 21.9%, DBA/2 3.1%. Mice were bred and housed under specific pathogen-free conditions at the Monash Medical Centre Animal Facilities (Clayton, Victoria, Australia) and used in experiments at 6-12 weeks of age. All animal studies were approved by the Monash University and Hudson Institute of Medical Research Monash Medical Centre B (MMCB) Animal Ethics Committees and conducted in accordance with the Australian Use Code (National Health and Medical Research Council of Australia).

### Verification of hPR3 expression in hPR3 knock-in mice

Expression of hPR3 in hPR3 knock-in mice was verified by RT-PCR and restriction digest. Total RNA was isolated from purified heterozygous mouse bone marrow neutrophils using TRIzol reagent (Life Technologies) and reversed-transcribed into cDNA using the Verso cDNA synthesis kit (Thermo Fisher Scientific). A homologous human and murine cDNA fragment was amplified by PCR using the same forward and backward primers DJ3536 (5’-AGCTACCCATCCCCCAAG-3) and DJ3538 (5’- ATGGCCAGGCACTGGGT-3) under the following conditions: 3 cycles at 58 °C annealing temperature, 3 cycles at 55 °C and 30 cycles at 52 °C. As a positive control β-actin was amplified with mouse β-actin primers (forward_5’- CTGGGCCGCCCTAGGCACCA-3’, reverse_ 5’-TGGCCTTAGGGTTCAGGG-3’) with an amplification program consisting of 3, 3 and 24 cycles with annealing temperatures at 65 °C, 62 °C and 59 °C, respectively. Differential transcription of the human and mouse PR3 cDNA was distinguished by SacI digestion of the PCR products, followed by agarose gel electrophoresis to determine fragment sizes.

### Assessment of hPR3 expression in hPR3 knock-in mice by western blot

Expression of hPR3 in hPR3 knock-in mice was assessed by western blot. In brief, freshly isolated bone marrow neutrophils (1 x 10^6^) from hPR3 knock-in mice were lysed in lysis buffer (50 mM Tris, 150 mM NaCl, 0.5 M EDTA, 0.5% deoxycholic acid, 0.1% SDS and 0.5% NP-40). Neutrophils isolated from peripheral blood of human and wild-type mice were used as controls. Cell lysates were prepared and resolved by 15% SDS-PAGE gel, followed by transfer of proteins onto a PVDF membrane by semidry electroblotting. After blocking with 5% non-fat dry milk blocker (Bio-Rad), membranes were incubated with a polyclonal primary antibody against both human and mouse PR3 (goat anti-mPR3, 1:1000) at 4°C overnight on a shaking incubator, followed by incubation with secondary antibodies (Mouse anti-goat-HRP, 1:1000) for 1 h at room temperature. Immunoreactive bands were detected by chemiluminescence substrate (SuperSignal West Pico Chemiluminescent Substrate, Pierce) and visualized with a ChemiDoc imaging system (Bio-Rad).

### Assessment of catalytic activity of hPR3 in hPR3 knock-in mice by FRET-based activity assay

Catalytic activity of hPR3 was assessed using a hPR3-specific FRET substrate (Abz- VADCADRQ-EDDnp). In brief, lysates of hPR3 knock-in mouse neutrophils (5 × 10^5^) were incubated with 5 µM substrate in activity assay buffer (50 mM Tris, 150 mM NaCl, 0.01% Triton X-100). Fluorescence was measured over time at λEx = 320 nm and λEm = 405 nm using a FLUOstar OPTIMA microplate reader. Recombinant human PR3 (rhPR3, 50 nM), recombinant mouse PR3 (rmPR3, 50 nM), lysates of human neutrophils (1 × 10^5^) and lysates of wild-type mouse neutrophils (5 × 10^5^) were included for comparison. Lysates of 5 × 10^5^ PR3/neutrophil elastase double knock-out neutrophils (*Prtn3/Elane*^-/-^) served as a negative control to exclude non-specific substrate cleavage.

### HLA-DR15^+^ patient cohort

Peripheral blood mononuclear cells were obtained from people with a confirmed diagnosis of PR3-AAV attending Monash Health (Melbourne, Victoria, Australia). Written informed consent was obtained from all participants, and all experiments were conducted in accordance with the guidelines of the Declaration of Helsinki. Human Research Ethics Committee Approval was obtained from the Monash Health Research Ethics Committee. Samples were stored at the Monash Vasculitis Biobank. Of 26 samples from people who were or had been PR3-ANCA^+^ that were HLA typed (Institute for Immunology and Infectious Diseases, Murdoch University, Perth,

Western Australia, Australia), 8 people expressed HLA-DRB1*15:01. Of these 8 HLA- DR15^+^ patients, five were PR3-ANCA^+^ at the time of sampling, were not additionally MPO-ANCA^+^ and had sufficient cells frozen for expansion. Three (A1-A3) had active disease, and two patients (R1 and R2) were in clinical remission but had a positive PR3-ANCA assay (table S6). All five patients were male. Four were of North-west European and one of Southern and Central Asian ethnicity.

### Synthetic peptides and assessment of T cell reactivity

To screen and validate CD4⁺ T cell epitopes of hPR3, IFN-γ and IL-17A ELISPOT assays were performed 10 days after immunization of DR15.hPR3^+^ mice with human PR3 (hPR3) overlapping peptides (either pooled or individual), recombinant hPR3, human complementary PR3 (cPR3), or mouse PR3 (mPR3), using a similar strategy to that used in previously published work (*11, 37*).

All test peptides were synthesized by GL Biochem (Shanghai, China) using Fmoc chemistry. Peptides were purified by RP-HPLC to at least 95% purity and lyophilised. Peptides were reconstituted in water to a concentration of 4-10 mg/ml, aliquoted and stored at -20 °C. For peptide immunization, mice were injected with either a pool of five peptides (10 μg each; total 50 μg/mouse) or with an individual peptide (20 μg/mouse). For protein immunization, mice received hPR3, cPR3 or mPR3 (100 μg/mouse). Peptides or proteins were prepared in 200 μL of saline/FCA (v/v=1:1) and injected subcutaneously at the base of the tail (100 μL per side).

Draining lymph nodes (dLNs) were collected 10 days after immunization. Lymph node cells were isolated and added to 96-well ELISPOT plates coated with unconjugated anti-IFN-γ (Mabtech AB, Sweden) or anti–IL-17A (BD Biosciences, San Jose, CA, USA) antibodies at a density of 1 × 10^6^ cells per well (or 5 × 10^5^ per well when additional restimulations were required). Cells were stimulated with peptide pools or individual peptides at a final concentration of 20 μg/mL and cultured for 18 h at 37 °C in 5% CO₂. Results were expressed as the number of IFN-γ or IL-17A spots per well.

### Fluorescence polarisation assay

The fluorescence polarisation assay was performed as previously described (*39*). Briefly, various concentrations ranging from 500 to 0 μM of PR3p43 (PR3_216-231_) test peptides were incubated in competition with 20 nM fluorescent reference peptides to bind to 100 nM recombinant HLA-DR15-StrepCLIP protein in the presence of 20 nM HLA-DM in Assay Buffer (100 mM trisodium citrate, pH 5.4, 50 mM NaCl, 5 mM EDTA). Fluorescent polarisation was measured after 24 hours of incubation at 37 °C using a PHERAstar microplate reader (BMG LABTECH). Peptide-binding curves were analysed by non-linear regression with Prism software (GraphPad Software version 10.2.0) using a sigmoidal dose-response curve. IC50 binding values were determined as the peptide concentration that inhibits 50% of reference peptide binding. Relative binding values (%) were calculated as the IC50 of the substituted peptide divided by the IC50 of the non-substituted peptide at the same concentration.

### Assessment of B cell reactivity

To assess the B cell response against PR3_216-231_, DR15.hPR3^+^ mice were immunized with 100 μg of PR3_216-231_ on days 0, 21, and 35. The first immunisation was prepared in saline/FCA (v/v = 1:1), followed by subsequent boosts in saline/FIA; v/v = 1:1). Experiments ended on day 42, and spleens and blood were collected for splenocyte isolation and serum preparation.

B cell responses were assessed by enumerating splenic B cells secreting anti-PR3_216- 231_ or anti-hPR3 antibodies via B cell ELISPOT, and by quantifying serum anti-PR3_216- 231_/hPR3 immunoglobulin (PR3-ANCA) levels via ELISA. PR3_216-231_ (20 μg/mL) and recombinant hPR3 (5 μg/mL) were used for cell restimulation in the ELISPOT assay and for plate coating in the ELISA. Control mice were immunized with 100 μg of OVA_323-339_. For both the ELISPOT (cell restimulation) and ELISA (plating coating), OVA_323-339_ (20 μg/mL) and OVA (5 μg/mL) were used.

For indirect immunofluorescence, human neutrophils were isolated from peripheral blood of healthy volunteers using density-gradient centrifugation. Isolated neutrophils were deposited onto microscope slides at a density of 20,000 cells per slide by cytocentrifugation. The cells were fixed with 4% PFA and acetone before blocking with 5% goat serum. Serum samples containing PR3-ANCA were applied to the fixed neutrophils for 30 min at room temperature. Bound autoantibodies were detected using FITC-conjugated goat anti-human IgG (F9512, Merck, Germany). Slides were then mounted with DAPI-containing antifade medium (ProLong™ Gold Antifade Mountant with DAPI, P36935; Thermo Fisher Scientific). PR3-ANCA staining patterns were visualized using a fluorescence microscope (Leica).

### Mass spectrometry analysis of HLA-DR15-presented hPR3 peptides

Splenocytes and lymph node cells isolated from HLA DR15.hPR3^+^ mice were co-incubated with heat-inactivated hPR3 protein at either 20 μg/mL or 100 μg/mL overnight prior to harvesting and snap-freezing cell pellets. Untreated spleen and lymph node cells were also harvested and snap frozen in parallel.

Peptide-HLA (pHLA) isolation using a pan-HLA-DR antibody (LB3.1), followed by peptide isolation via RP-HPLC fractionation, was performed as described in Purcell *et al.* (*53*). In brief, cell pellets were pulverised in a cryomill and pHLA complexes solubilised in a non-denaturing lysis buffer containing 0.5% IGEPAL. Intact pHLA complexes were subsequently immunopurified with LB3.1 cross-linked protein A sepharose resin, eluted with 10% glacial acetic acid, and the extracted peptide ligands fractionated using RP-HPLC. Peptide containing fractions were vacuum-concentrated and resuspended in 100mM triethylammonium bicarbonate. Resuspended peptides were reduced with 10mM tris(2-carboxyethyl)phosphine, alkylated with 20mM of iodoacetamide, and acidified to a final concentration of 1% formic acid (FA).

Peptide samples were loaded onto PureTips (Evosep Biosystems) according to the manufacturer’s instructions and analysed on a Bruker TimsTOF Pro2 mass spectrometer coupled to an Evosep One liquid chromatography system. Separation was performed on an IonOpticks Aurora Elite column (15 cm × 75 μm × 1.7 μm, 120 Å pore size) using the Zoom Whisper 20 SPD method as described (*54, 55*) Samples were acquired using a data-dependent acquisition method with the following parameters: m/z range 100-1700, capillary voltage 1600 V, and a target intensity of 30,000. Ion mobility was enabled with a TIMS ramp of 0.60 to 1.60 Vs/cm^2^ over 166 ms.

Acquired data files were searched using the Peaks Online software (v11, Bioinformatics Solutions Inc.) against the reviewed *Mus Musculus* proteome appended with the human PR3 protein sequence downloaded on 07/06/2024 from Uniprot. The database search parameters used were precursor mass error tolerance of 20 ppm, fragment mass error tolerance 0.02 Da, and no enzyme specificity. Carbamidomethylation of cysteines was set as a fixed modification. Deamidation of asparagine and glutamine, and oxidation of methionine, were set as variable modifications. All datasets were analysed at a 5% peptide false discovery rate cut off.

### HLA-DRA1*01:01/DRB1*15:01- PR3_216-231_ purification and tetramer production

To enhance water solubility, two glutamic acid residues were added at the C-terminus of the synthesised PR3_216-_231-Cys (IDSFVIWGCATRLFPD*EE*) (extra residues in *italics*), and to PR3_216-231_-Abu (IDSFVIWG<u>X</u>ATRLFPD*EE*) (mutated residue underlined and extra residues in *italics*) peptides. The peptide loading process followed the method described in a previous study (*28*). Briefly, HLA-DR15-StrepCLIP was digested with Factor Xa (New England BioLabs, MA, USA) to cleave the covalently linked Strep-TactinII-CLIP peptide at room temperature (RT) for 6 hours. A 20-fold molar excess of PR3_216-231_-Cys or PR3_216-231_-Abu peptide was then loaded onto the StrepClip-Factor Xa cleaved HLA-DR15 in a buffer composed of 50 mM trisodium citrate pH 5.4, and 5 mM EDTA at 37 °C for 24 hours, with HLA-DM serving as a catalyst. The peptide-loaded HLA-DR15 was separated from the unloaded HLA- DR15-StrepCLIP using Strep-Tactin Sepharose (IBA, Göttingen, Germany). HLA- DR15-PR3_216-231_-Cys/-PR3_216-231_-Abu complexes were buffer exchanged into 10 mM Tris ph 8.0, 150 mM NaCl, then biotinylated using E. coli BirA, and purified with a HiTrapTM Desalting Column (Cytiva). Tetramers were produced by conjugating biotinylated HLA-DR15-peptide monomers with streptavidin-PE at a 4:1 molar ratio. The fluorochrome was added in ten steps, with incubation for 10 minutes at RT in the dark during each addition.

### Crystallization and structure determination

Before crystallisation trials, the HLA-DRA1*01:01/DRB1*15:01-PR3_216-231_-Abu/- PR3_216-231_-Cys complexes were subjected to HRV 3C protease cleavage at 4 °C for 24 hours to remove the fos/jun leucine zippers, followed by purification through anion exchange chromatography (HiTrap Q HP column: Cytiva). The target fractions were pooled and buffer exchanged into 10 mM Tris pH 8.0, 150 mM NaCl. The solution at a concentration of 10 mg/ml was used for broad screening crystallisation trials at the Monash Molecular Crystallisation Platform using an automated robotic NT8 system. Conditions that produced crystals were further scaled up and optimised via the hanging-drop vapour diffusion method in 24-well plates. The complex HLA-DR15- PR3_216-231_-Abu/ -PR3_216-231_-Cys crystallised in 8% v/v Tacsimate and 20% w/v PEG3350. Crystals were cryoprotected in the mother liquor well solution supplemented with 20% glycerol before flash freezing in liquid N2. Diffraction data were collected on the MX2 Beamline of the Australian Synchrotron, using an Eiger X16 M detector and processed and scaled with XDS and CCP4 Software Suite, version 8.0. HLA-DRA1*01:01/DRB1*15:01-PR3_216-231_-Abu/-PR3_216-231_-Cys binaries were solved by molecular replacement in PHASER (CCP4 Software Suite, version 8.0) using search model for HLA-DR4 (PDB ID: 9NIH) (*30*). PyMol version 2.1 (http://www.pymol.org/) was used to generate all structural figures.

### Enrichment of antigen-specific CD4^+^ T cells *in vitro*

For the enrichment of PR3_216-231_-specific mouse CD4^+^ T cells, CD4^+^ T cells were isolated from the dLNs and spleens of PR3_216-231_-Cys-immunized DR15.hPR3 mice using a mouse CD4^+^ T cell isolation kit (Miltenyi Biotec). Expansion of PR3_216-231_- specific human CD4^+^ T cells was performed using peripheral blood mononuclear cells (PBMCs) from five HLA-DR15^+^ PR3-ANCA^+^ patients (R1, R2, A1, A2, and A3). For each individual patient, PBMCs were co-cultured with mitomycin C-treated (Accord) PR3_216-231_-Cys- or PR3_216-231_-Abu-pulsed antigen presenting cells (APCs) at a ratio of 1:10 (APC:T) in T cell culture media [RPMI supplemented with 10% fetal calf serum, 0.5 mg/mL penicillin/streptomycin, 2 mM GlutaMAX-I, 1 mM sodium pyruvate, 55 μM 2-mercaptoethanol, and recombinant human IL-2 (10 U/mL for mouse T cells or 25 U/mL for human T cells; PeproTech)] at 37 °C with 5% CO_2_. hPR3.DR15^+^ mouse BMDCs and the DR15^+^ human monocytic leukemia cell line THP-1 were used as APCs for the expansion of mouse and human PR3_216-231-_specific CD4^+^ T cells, respectively. Cells received 1-3 restimulations with PR3_216-231_-pulsed APCs at 10-14- day intervals. Alternatively, PBMCs from patients R2, A1, and A3 were labelled with 1 μM CellTrace Violet (CTV) and co-cultured individually for 7 days with mitomycin C- treated PR3_216-231_-Cys- or PR3_216-231_-Abu-pulsed APCs (APC:T = 1:10) under the same conditions described above. PBMCs from patients R1 and A2 were excluded from CTV labelling due to insufficient cell numbers for this experiment.

### Cell labelling, tetramer and antibody staining

Identification of PR3_216-231_-specific CD4^+^ T cells was performed using PE-conjugated HLA-II tetramers. Cells were stained with PE-labelled HLA-DRA1*01:01/DRB1*15:01 (DR15)-PR3_216-231_ tetramers for 1 hour at room temperature, washed, and subsequently stained with a cocktail of fluorescently conjugated antibodies for 30 minutes at 4 ℃ to identify epitope-specific cells among mouse CD4^+^ T cells (CD45, CD3, CD4, CD8, Fixable Viability Stain 700) and human CD4^+^ T cells (CD3, CD4, CD8, LIVE/DEAD Fixable Near-IR) using the gating strategy in Fig. S10, A to C. Cells were analyzed on an LSRFortessa X-20 or sorted on a FACSAria Fusion cell sorter (BD Biosciences). The antibodies and reagents used are listed in Table S8.

### Analysis of epitope-specific T cell repertoires

Mouse DR15-PR3_216-231_ tetramer^+^ cells were single-cell sorted from *in vitro* expanded dLN and splenic CD4^+^ T cells. Individual human DR15-PR3_216-231_ tetramer^+^ cells were sorted from either *in vitro* expanded (R1, R2, A1, A2, and A3) or CTV-labelled (R2, A1, and A3) CD4^+^ T cell populations. For both mouse and human samples, mRNA from single cells was reverse-transcribed, and a multiplexed nested polymerase chain reaction (PCR) strategy was employed to amplify cDNA encoding the CDR3α and CDR3β regions using a panel of Vα- and Vβ-specific oligonucleotide primers (*33, 56*). PCR products were purified and sequenced using an ABI 3730 DNA Sequencer, and *TRBV* and *TRAV* gene usage was determined via the IMGT/V-QUEST database (www.imgt.org/IMGT_vquest) (*57*). TCR clonotypes and similarity were analysed using the TCRdist pipeline as described unless otherwise indicated below (*33*). Kernel principal component analysis (kPCA) and nearest neighbour (NN) distance analysis were performed using the TCRdist matrix generated by the default pipeline. Aesthetic modifications to the original figure scripts include showing clone size by the size of the circles for the kPCA plots and the addition of bar histograms (bin size 10 for individual chains, 20 for paired chains) to the NN distance graphs. To enable comparison between human and mouse TCRs, a pairwise CDR3-only distance was derived and plotted as a heatmap (by removing the weighing of CDR1 and CDR2 in the original TCRdist method) to remove the effect of different germ-line sequences between species. Dendrograms show hierarchical clustering of TCRs within each repertoire using UPGMA (average linkage) on the pairwise CDR3 distance matrix.

### In vitro TCR expression and tetramer staining

TCR expression in vitro as described in previous studies (*28, 40*). Genes encoding TCR α- and β-chains (Genscript) were cloned into the lentivirus vectors (Biosettia) pLV-EF1α-MCS-IRES-GFP and pLV-EF1α-MCS-IRES-RFP, respectively, to generate pLV-EF1α-TCRα-IRES-GFP and pLV-EF1α-TCRβ-IRES-RFP. A mixture containing a total of 1 μg (1:1 ratio) of pLV-EF1α-TCRα-IRES-GFP and pLV-EF1α-TCRβ-IRES-RFP plasmids, together with 500 ng of pLV-CD3γ,δ,ε, and ζ sub-units and 5 μl Fugene6 (Promega), was incubated in 171 μl OPTI-MEM I (1x) (Thermo Fisher Scientific) media at RT for 15 min. Approximately, 0.7×10^6^ HEK293T cells were transfected with the Fugene-DNA mixture and cultured in 3 ml of fresh RF10 media: RPMI-1640 (Thermo Fisher Scientific) supplemented with 50 IU ml−1 penicillin, 50 μg ml−1 streptomycin, 2 mM glutamine, 1× nonessential amino acids, 1 mM pyruvate, 10 mM HEPES (all from Thermo Fisher Scientific), 50 μM 2- mercaptoethanol (Merck), and 10% FCS (Merck) in a 6-well plate. The transfected HEK293T cells were incubated at 37 °C for 24 hours before pMHC tetramer staining. Transfected 293T cells were washed twice with FACS buffer (Phosphate buffered saline, 10% Fetal Calf Serum (FCS; Merck)) before being stained with 1 μg of HLA- HLA-DR15-PR3_216-231_Cys tetramer conjugated with PE for 1 hour at RT, followed by 1 hour staining with a 1:100 dilution of BUV395 mouse anti-human CD3 (clone UCHT1, BD Biosciences) on ice in the dark. The transfected HEK293T cells were washed 4 times with FACS buffer before being live/dead-stained with DAPI (BD Biosciences). The cells were analysed by flow cytometry (LSR II; BD Biosciences; BD FACSDiva- 8.0.1 software). The CD3^+^ and HLA-DR15-PR3_216-231_Cys tetramer-positive cells were determined using Flowjo v10.6.0. Two independent experiments were conducted in duplicate.

### Assessment of anti-PR3 autoimmune glomerulonephritis

DR15.hPR3^+^ mice were immunized subcutaneously with 100 μg of PR3_216-231_ on days 0 and 7, first in 0.9% saline/FCA (v/v=1:1) and then in 0.9% saline/Freund’s incomplete adjuvant (FIA). On day 17, PR3 was deposited in glomeruli by recruiting neutrophils using a low dose of intravenously injected heterologous anti-mouse basement membrane antibodies (*58, 59*). Experiments were concluded on day 21.

Urine was collected over an 18-hour period using metabolic cages prior to euthanasia. Blood was collected on the day of euthanasia to obtain serum samples. Albuminuria and serum creatinine levels were measured using enzyme-linked immunosorbent assays (ELISAs). Glomerular abnormalities were assessed on Periodic acid-Schiff-stained, 3-μm-thick, paraffin-embedded kidney sections mounted on coded slides. The percentage of abnormal glomeruli was determined by examining a minimum of 50 glomeruli per mouse. Abnormalities included glomerular hypercellularity, capillary wall thickening and segmental fibrinoid necrosis. CD4⁺ T cells, macrophages, and neutrophils were identified by immunoperoxidase staining of 4-μm-thick, periodate-lysine–paraformaldehyde (PLP)-fixed, frozen kidney sections. Primary monoclonal antibodies used were rat anti-mouse CD4 (clone GK1.5, BioXcell) for CD4⁺ T cells, rat anti-mouse CD68 (clone FA/11, BioXcell) for macrophages, and rat anti-mouse Gr-1 (clone RB6-8C5, BioXcell) for neutrophils. Cells were visualized using a VECTASTAIN® Elite ABC Kit (Vector Laboratories) and 3,3′-diaminobenzidine (DAB) substrate (Sigma-Aldrich), with rabbit anti-rat biotin (Vector Laboratories) as the secondary antibody. For quantification, at least 50 glomeruli and 20 consecutive high-power (40× objective) interstitial fields, excluding perivascular regions, were assessed per mouse. Results were expressed as the number of cells per glomerular cross-section (cells/gcs) and as total cell counts across the 20 interstitial fields.

### Cell isolation and Flow cytometry

Kidney single-cell suspensions were prepared by mincing and digesting the kidneys in 1 mL of Hanks’ Balanced Salt Solution (HBSS; Gibco) containing 4 mg/mL Collagenase D and 100 μg/mL DNase I (Merck) for 20 minutes at 37 °C with constant shaking. The digested tissue was then filtered through 40 μm strainers, and red blood cells were removed using red blood cell lysis buffer.

Cell surface staining of single-cell suspensions was performed using fluorescently conjugated antibodies following FcγRIII/II receptor blocking with anti-CD16/32. Intracellular staining was conducted after fixation and permeabilization using the FoxP3/Transcription Factor Staining Buffer Set (eBioscience). Dead cells were excluded using the Live/Dead marker FVS700. All samples were acquired on a 5-laser spectral flow cytometer (Aurora, Cytek Biosciences) and analysed using FlowJo software v10.10 (BD Biosciences).

After down-sampling to 20,000 CD45.2⁺ cells per sample using the DownSample plugin and concatenating the datasets with the concatenation tool, Uniform Manifold Approximation and Projection (UMAP) analysis was performed using the FlowJo plugin with standard settings to visualize cells and clusters. Clustering parameters included CD11b, Ly6G, F4/80, CD11c, Ly6C, TCRβ, CD4, CD8, FoxP3, CD44, CD62L, CD69, and CD19. The antibodies used are listed in table S8.

## Supplementary Materials

Figs. S1 to S14 Tables S1 to S8

## References and Notes

## Acknowledgments

The authors acknowledge the staff and the use of equipment at the Monash Health Translation Platform (MHTP) FlowCore Facility, the MHTP Animal Facility and the MHTP Histology Platform. The authors thank Reinhard Fässler and Therese Dau for assistance with the hPR3 mouse.

## Author contributions

Conceptualization: ARK, JRo, HHR, CL and LRS; funding acquisition: ARK, DJ, JRo, HHR, AWP and NLL; Mouse generation and characterization: LA, MM, DJ and LF; Human samples and clinical data: ARK, LRS and JRy; Investigation: CL, LRS, JJL, TJL, SG, CMJ, LH, PP and MTT; Data Analysis: CL, LRS, JJL, TJL, SG, CMJ, LA, JBZ, MTT, DC, DJ, HHR, JRo and ARK; Writing -original draft: CL, LS, JJL, TJL, JBZ, MTT, JRo and ARK; Writing - review and editing: CL, LRS, JJL, TJLK, SG, CJ, JBZ, LA, PP, MTT, DC, JRy, LF, MM, DJ, HHR, NLL, AWP, JRo and ARK.

## Data and materials availability

The mass spectrometry proteomics data have been deposited to the ProteomeXchange Consortium via the PRIDE partner repository (*60*) with a dataset identifier that will be released on full publication. The structures were deposited in the PDB database DR15-PR3-p43-Cys, PDB ID: 43TP and DR15-PR3- p43-Abu, PDB ID: 43TO.

---

**Fig. S1. Experimental design for the generation of hPR3 knock-in mice.** In the wild-type genome, exons 2-5 of the murine PR3 (*Prtn3*) were replaced with the human PR3 (*PRTN3*) gene, while the murine promoter region up to exon 1, containing only the murine signal peptide, was retained to ensure *PRTN3* expression under physiological regulatory control. A neomycin resistance cassette (NEO) flanked by two loxP sites at the 3′ end of the *PRTN3* gene segment was included in the targeting vector for selection of recombinant ES cells, and subsequently excised by crossing the knock-in mice with a Cre deleter line on a C57BL/6J background. by Cre recombinase in ES cells showing homologous replacement of the four murine exons.

**Fig. S2. Human PR3 expression and catalytic activity in hPR3 knock-in mice.** (**A**) RNA was isolated from neutrophils of heterozygous hPR3 knock-in mice, reverse-transcribed into cDNA and a homologous 440-bp fragment amplified by PCR. Differential transcription of human and mouse PR3 was distinguished by SacI digestion that selectively cleaves the human PR3 fragment into products of 270 bp and 170 bp, while the mouse fragment remained undigested (M, molecular weight maker). (**B**) Peripheral blood neutrophils (1x10^6^) from human, wild-type control mice (WT) and hPR3 knock-in (hPR3^+/+^) mice were lysed and analyzed by SDS PAGE and immunoblotting. Human and mouse PR3 were detected by a goat anti-mPR3 polyclonal antibody and expression levels determined semi-quantitatively compared to increasing amounts of recombinant (r)hPR3. Results are representative of one of three independent experiments, presented as mean±SEM (n=3). (**C**) hPR3 catalytic activity of hPR3^+/+^ mouse neutrophils was assessed with a PR3-specific FRET substrate (Abz-VADCADRQ-EDDnp, λEx=320 nm, λEm=405 nm). hPR3^+/+^ neutrophil (5×10^5^) lysates were incubated with 5 µM substrate. Controls included rhPR3 (50 nM), rmPR3 (50 nM), and human (1×10^5^) and WT mouse (5×10^5^) neutrophil lysates. PR3/NE knock-out neutrophil (PR3/NE^-/-^, 5×10^5^) lysates served as a negative control, excluding non-specific substrate cleavage. Data are presented as mean±SEM (n=3).

**Fig. S3. Lack of reactivity to peptides derived from murine and human PR3 exon 1.** Antigen-specific T cell responses were assessed by IL-17A and IFN-γ ELISPOT assays 10 days after immunization with mouse (mPR3_1-16_, mPR3_5-21_, mPR3_10-26_) or human (hPR3_1-16_, hPR3_5-21_) PR3 exon 1-derived peptides. IL-17A and IFN-γ production was measured following *ex vivo* restimulation of draining lymph node (inguinal and para-aortic) cells with individual peptides spanning mouse or human PR3 exon 1. Horizontal bars indicate means for each group (n = 8 per group).

**Fig. S4. Lack of consistent reactivity to complementary hPR3 (cPR3) peptides.** (**A**) The complementary amino acid sequence of hPR3 exons 2-5. The position of the 31 cPR3 peptides included in the cPR3 peptide library and screening are shown. Numbering begins at methionine (M) at amino acid position 185, as most top-ranked IEDB-predicted epitopes fall within exons 2-4. The remaining predicted epitopes localize to amino acids 226-243 in exon 5 of cPR3 and were excluded from screening due to high sequence identity with PR3p45 (PR3_226-241_). (**B** to **D**) Three independent experiments assessing antigen-specific T cell responses to 31 cPR3 peptides by IL- 17A and IFN-γ ELISPOT assays 10 days after immunization of hPR3.DR15^+^ mice with peptide pools 1-3 (pool 1, peptides 1-10; pool 2, peptides 11-20, pool 3, peptides 21- 31). IL-17A and IFN-γ production was measured followed *ex vivo* restimulation of draining lymph node cells with the corresponding peptide sub-pools (3 to 4 peptides per pool). Horizontal bars indicate means for each group (n = 4 to 5 per group).

**Fig. S5. hPR3 peptide 43 (PR3_216-231_) is confirmed as the immunodominant epitope, whereas peptide 47 (PR3_236-251_) is non-immunogenic.** (**A** and **B**) Quantification of antigen-specific T cell responses to the corresponding individual hPR3 peptides 10 days after immunization of hPR3.DR15^+^ mice with a peptide pool consisting of hPR3 peptides 41-45 (41/PR3_206-221_, 42/PR3_211-226_, 43/PR3_216-231_, 44/PR3_221-236_, 45/PR3_226-241_) (**A**), or to hPR3 peptides 17 (PR3_86-101_, non-immunogenic control), 43 (PR3_216-231_) and 47 (PR3_236-251_) 10 days after immunization with recombinant hPR3 (rhPR3) or hPR3 peptide 43 (PR3_216-231_) (**B**). Horizontal bars indicate means for each group (n = 2 to 8 per group).

**Fig. S6. PR3_216-231_ induces autoreactive B cell response.** (**A** and **B**) Quantification of autoreactive B cell response 42 days after immunization with peptides (PR3_216-231_ or control peptide OVA_323-339_) or proteins (hPR3 or OVA) on day 0, with boosts on days 21 and 35. Antibody production against the corresponding peptides (**A**) or proteins (**B**) was measured by B cell ELISPOT (left) and ELISA (right). Horizontal bars indicate means for each group (n = 10 per group). Statistical analysis was performed using Student’s *t* test. \*\*\**P* < 0.001; \*\*\*\**P* < 0.0001.

**Fig. S7. Antibodies from the sera of PR3_216-231_ immunized hPR3.DR15^+^ mice do not induce a cANCA pattern in human neutrophils.** Neutrophils isolated from peripheral blood of healthy doors were incubated with serum samples containing PR3- ANCA, followed by detection of bound antibodies using FITC-conjugated anti-human IgG. Representative images show PR3-ANCA staining by indirect immunofluorescence.

**Fig. S8. HLA-DR15-presented peptide derived from the hPR3_103-119_ region is non-immunogenic.** (**A** and **B**) Quantification of antigen-specific T cell responses to individual hPR3 peptides 10 days after immunization of hPR3.DR15^+^ mice with hPR3 protein (**A**) or with individual hPR3 peptides (**B**). The peptides tested were PR3p20/PR3_101-116_ and PR3p21/PR3_106-121_. Responses were measured by IFN-γ ELISPOT assays. PR3p40/PR3_201-216_ and PR3p43/PR3_216-231_ were used as non-immunogenic and immunogenic control peptides, respectively.

**Fig. S9. Superposition of HLA-DR15 presenting PR3_216-231_Abu and PR3_216-231_Cys.** The peptides are depicted as sticks and coloured lemon (PR3_216-231_Abu) and cyan (PR3_216-231_Cys). Amino acid residues are labelled using single-letter codes.

**Fig. S10. Flow cytometric gating strategies for single-cell sorting of DR15- PR3_216-231_-tetramer^+^ CD4^+^ T cells.** Representative gating strategies used to identify DR15-PR3_216-231_-tetramer^+^ cells among *in vitro*-expanded CD4^+^ T cells from the draining lymph nodes and spleens of PR3_216-231_-immunized hPR3.DR15^+^ mice (**A**), *in vitro*-expanded CD4^+^ T cells from DR15^+^ PR3-AAV patient PBMCs (**B**), and CTV-labelled and *in vitro*-expanded CD4^+^ T cells from the same patient PBMCs (**C**).

**Fig. S11. DR15-PR3_216-231_-specific CD4^+^ TCR repertoires in humanized hPR3.DR15^+^ mice and DR15^+^ PR3-AAV patients.** (**A** and **B**) V-J gene pairing of each clonotype in the PR3_216-231_-specific CD4^+^ TCR repertoires in humanized hPR3.DR15^+^ mice (**A**) and DR15^+^ PR3-AAV patients (**B**). The vertical height of each gene segment is proportional to its frequency within the repertoire. V-J pairing between and across α and β chains are shown as curved lines joining each gene. Arrows denote fold change (fc) of gene usage relative to the background human TCR dataset in the TCRdist pipeline. One arrowhead, 2 < fc < 4; two arrowheads, 4 < fc < 8; five arrowheads, fc > 32.

**Fig. S12. HLA-DR15-PR3_216-231_ tetramer staining of cell line expressing TRAV 16_TRBV19 TCR.** HEK293 T cells transiently co-transfected with a PR3_216-231_-specific hPR3.DR15^+^ mouse TRAV16-TRBV19-using TCR and CD3γδεζ were stained with either HLA-DR15-CLIP (left) or HLA-DR15-PxR3_216-231_ tetramer (right).

**Fig. S13. DR15-PR3_216-231_-specific CD4^+^ TCR clonotypes and their expansion in individual DR15^+^ PR3-AAV patients**. kPCA projection of the TCRdist matrix for each individual DR15^+^ PR3-AAV patients with active disease (top row) and in remission (bottom row). TCR clones with clonal exxpansion are shown as circles with size corresponding to clone size. Colours are based on V or J gene frequency within the repertoire as follows: red (most frequent), green, blue, cyan, magenta followed by assorted colours for rare frequencies.

**Fig. S14. UMAP analysis of kidney leukocytes.** Markers including CD11b, Ly6G, F4/80, CD11c, Ly6C, TCRβ, CD4, CD44, CD62L and Foxp3 were used to identify neutrophils, macrophages subsets, and CD4^+^ T cell subsets among kidney leukocytes UMAP analysis was performed, and the heatmap indicates the relative expression levels of these markers.

