## Supplementary Figures 1-14 for "Unconventional presentation of an immunodominant HLA-DR15-restricted nephritogenic proteinase 3 epitope implicated in vasculitis"

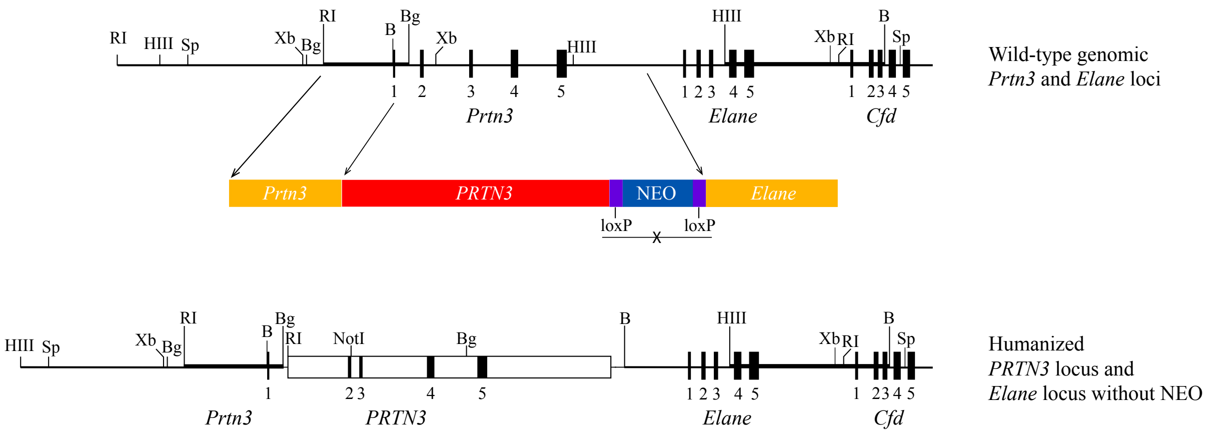


**Fig. S1. Experimental design for the generation of hPR3 knock-in mice.**

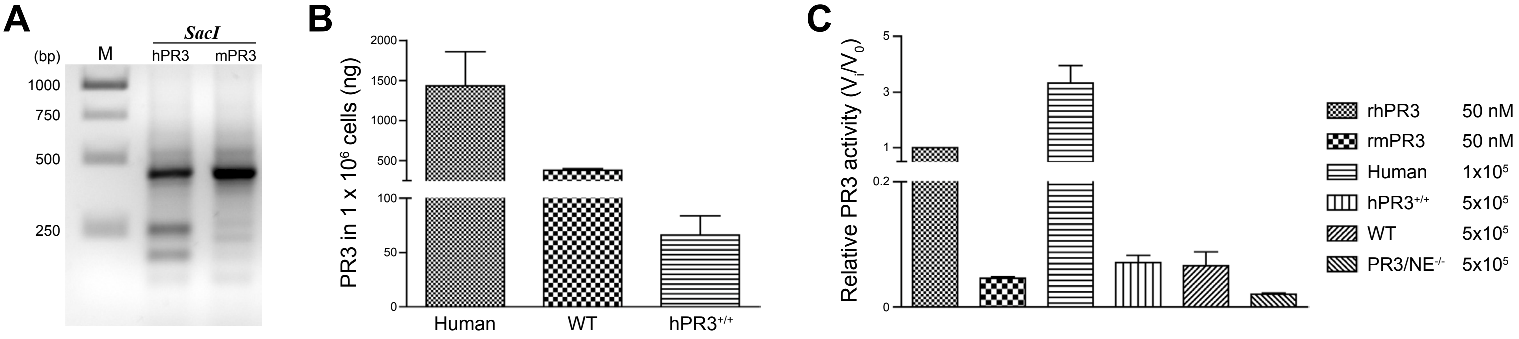


**Fig. S2. Human PR3 expression and catalytic activity in hPR3 knock-in mice.**

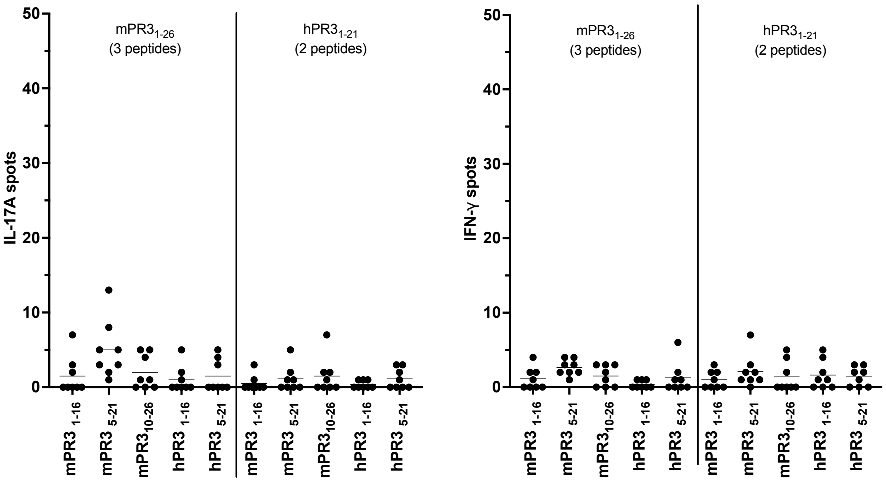


**Fig. S3. Lack of reactivity to peptides derived from murine and human PR3 exon 1.**

**A**

| EXON 1 | Amino acid | 1 | 2 | 3 | 4 | 5 | 6 | 7 | 8 | 9 | 10 | 11 | 12 | 13 | 14 | 15 | 16 | 17 | 18 | 19 | 20 |
|---|---|---|---|---|---|---|---|---|---|---|---|---|---|---|---|---|---|---|---|---|---|
| | Sense | M | A | H | R | P | P | S | P | A | L | A | S | V | L | L | A | L | L | L | S |
| | *Complementary* | *H* | *S* | *V* | *P* | *G* | *G* | *A* | *R* | *G* | *Q* | *R* | *G* | *H* | *Q* | *Q* | *G* | *Q* | *Q* | *Q* | *A* |

| EXON 2 | Amino acid | 21 | 22 | 23 | 24 | 25 | 26 | 27 | 28 | 29 | 30 | 31 | 32 | 33 | 34 | 35 | 36 | 37 | 38 | 39 | 40 | 41 | 42 | 43 | 44 | 45 | 46 | 47 | 48 | 49 | 50 |
|---|---|---|---|---|---|---|---|---|---|---|---|---|---|---|---|---|---|---|---|---|---|---|---|---|---|---|---|---|---|---|---|
| | Sense | G | A | A | R | A | A | E | I | V | G | G | H | E | A | Q | P | H | S | R | P | Y | M | A | S | L | Q | M | R | G | N |
| | *Complementary* | *T* | *S* | *G* | *S* | *S* | *R* | *L* | *D* | *H* | *A* | *P* | *V* | *L* | *R* | *L* | *W* | *V* | *G* | *P* | *G* | *V* | *H* | *G* | *G* | *Q* | *L* | *H* | *P* | *P* | *V* |
| | *Peptide* | | | | | | | | | | | | | | | *31* | | | | | *30* | | | | | *29* | | | | | *28* |

| | Amino acid | 51 | 52 | 53 | 54 | 55 | 56 | 57 | 58 | 59 | 60 | 61 | 62 | 63 | 64 | 65 | 66 | 67 | 68 | 69 | 70 | 71 | 72 | 73 | 74 | 75 | 76 | 77 | 78 | 79 | 80 |
|---|---|---|---|---|---|---|---|---|---|---|---|---|---|---|---|---|---|---|---|---|---|---|---|---|---|---|---|---|---|---|---|
| | Sense | P | G | S | H | F | C | G | G | T | L | I | H | P | S | F | V | L | T | A | A | H | C | L | R | D | I | P | Q | R | L |
| | *Complementary* | *R* | *A* | *A* | *V* | *E* | *A* | *S* | *A* | *G* | *Q* | *D* | *V* | *G* | *A* | *E* | *H* | *Q* | *R* | *G* | *R* | *V* | *A* | *Q* | *P* | *V* | *Y* | *G* | *L* | *A* | *Q* |
| | *Peptide* | | | | | *27* | | | | | *26* | | | | | *25* | | | | | *24* | | | | | *23* | | | | | *22* |

| EXON 3 | Amino acid | 81 | 82 | 83 | 84 | 85 | 86 | 87 | 88 | 89 | 90 | 91 | 92 | 93 | 94 | 95 | 96 | 97 | 98 | 99 | 100 | 101 | 102 | 103 | 104 | 105 | 106 | 107 | 108 | 109 | 110 |
|---|---|---|---|---|---|---|---|---|---|---|---|---|---|---|---|---|---|---|---|---|---|---|---|---|---|---|---|---|---|---|---|
| | Sense | V | N | V | V | L | G | A | H | N | V | R | T | Q | E | P | T | Q | Q | H | F | S | V | A | Q | V | F | L | N | N | Y |
| | *Complementary* | *H* | *V* | *H* | *H* | *E* | *S* | *G* | *V* | *V* | *H* | *P* | *R* | *L* | *L* | *G* | *G* | *L* | *L* | *V* | *E* | *R* | *H* | *S* | *L* | *H* | *K* | *Q* | *V* | *V* | *V* |
| | *Peptide* | | | | | *21* | | | | | *20* | | | | | *19* | | | | | *18* | | | | | *17* | | | | | *16* |

| | Amino acid | 111 | 112 | 113 | 114 | 115 | 116 | 117 | 118 | 119 | 120 | 121 | 122 | 123 | 124 | 125 | 126 | 127 | 128 | 129 | 130 | 131 | 132 | 133 | 134 | 135 | 136 | 137 | 138 | 139 | 140 |
|---|---|---|---|---|---|---|---|---|---|---|---|---|---|---|---|---|---|---|---|---|---|---|---|---|---|---|---|---|---|---|---|
| | Sense | D | A | E | N | K | L | N | D | V | L | L | I | Q | L | S | S | P | A | N | L | S | A | S | V | A | T | V | Q | L | P |
| | *Complementary* | *V* | *R* | *L* | *V* | *F* | *Q* | *V* | *V* | *N* | *E* | *E* | *D* | *L* | *Q* | *A* | *A* | *W* | *G* | *V* | *E* | *T* | *G* | *G* | *D* | *G* | *C* | *D* | *L* | *Q* | *W* |
| | *Peptide* | | | | | *15* | | | | | *14* | | | | | *13* | | | | | *12* | | | | | *11* | | | | | *10* |

| EXON 4 | Amino acid | 141 | 142 | 143 | 144 | 145 | 146 | 147 | 148 | 149 | 150 | 151 | 152 | 153 | 154 | 155 | 156 | 157 | 158 | 159 | 160 | 161 | 162 | 163 | 164 | 165 | 166 | 167 | 168 | 169 | 170 |
|---|---|---|---|---|---|---|---|---|---|---|---|---|---|---|---|---|---|---|---|---|---|---|---|---|---|---|---|---|---|---|---|
| | Sense | Q | Q | D | Q | P | V | P | H | G | T | Q | C | L | A | M | G | W | G | R | V | G | A | H | D | P | P | A | Q | V | L |
| | *Complementary* | *L* | *L* | *V* | *L* | *W* | *H* | *G* | *V* | *A* | *G* | *L* | *A* | *Q* | *G* | *H* | *A* | *P* | *A* | *A* | *H* | *T* | *G* | *V* | *V* | *G* | *W* | *G* | *L* | *D* | *Q* |
| | *Peptide* | | | | | *9* | | | | | *8* | | | | | *7* | | | | | *6* | | | | | *5* | | | | | *4* |

| | Amino acid | 171 | 172 | 173 | 174 | 175 | 176 | 177 | 178 | 179 | 180 | 181 | 182 | 183 | 184 | 185 | 186 | 187 | 188 | 189 | 190 | 191 | 192 | 193 | 194 | 195 | 196 | 197 | 198 | 199 | 200 |
|---|---|---|---|---|---|---|---|---|---|---|---|---|---|---|---|---|---|---|---|---|---|---|---|---|---|---|---|---|---|---|---|
| | Sense | Q | E | L | N | V | T | V | V | T | F | F | C | R | P | H | N | I | C | T | F | V | P | R | R | K | A | G | I | C | F |
| | *Complementary* | *L* | *L* | *E* | *I* | *D* | *G* | *H* | *D* | *G* | *E* | *E* | *A* | *P* | *W* | *M* | <--- | | | | | | | | | | | | | | |
| | *Peptide* | | | | | *3* | | | | | *2* | | | | | *1* | | | | | | | | | | | | | | | |

| EXON 5 | Amino acid | 201 | 202 | 203 | 204 | 205 | 206 | 207 | 208 | 209 | 210 | 211 | 212 | 213 | 214 | 215 | 216 | 217 | 218 | 219 | 220 | 221 | 222 | 223 | 224 | 225 | 226 | 227 | 228 | 229 | 230 |
|---|---|---|---|---|---|---|---|---|---|---|---|---|---|---|---|---|---|---|---|---|---|---|---|---|---|---|---|---|---|---|---|
| | Sense | G | D | S | G | G | P | L | I | C | D | G | I | I | Q | G | I | D | S | F | V | I | W | G | C | A | T | R | L | F | P |

| | Amino acid | 231 | 232 | 233 | 234 | 235 | 236 | 237 | 238 | 239 | 240 | 241 | 242 | 243 | 244 | 245 | 246 | 247 | 248 | 249 | 250 | 251 | 252 | 253 | 254 | 255 | 256 |
|---|---|---|---|---|---|---|---|---|---|---|---|---|---|---|---|---|---|---|---|---|---|---|---|---|---|---|---|
| | Sense | D | F | F | T | R | V | A | L | Y | V | D | W | I | R | S | T | L | R | R | V | E | A | K | G | R | P |

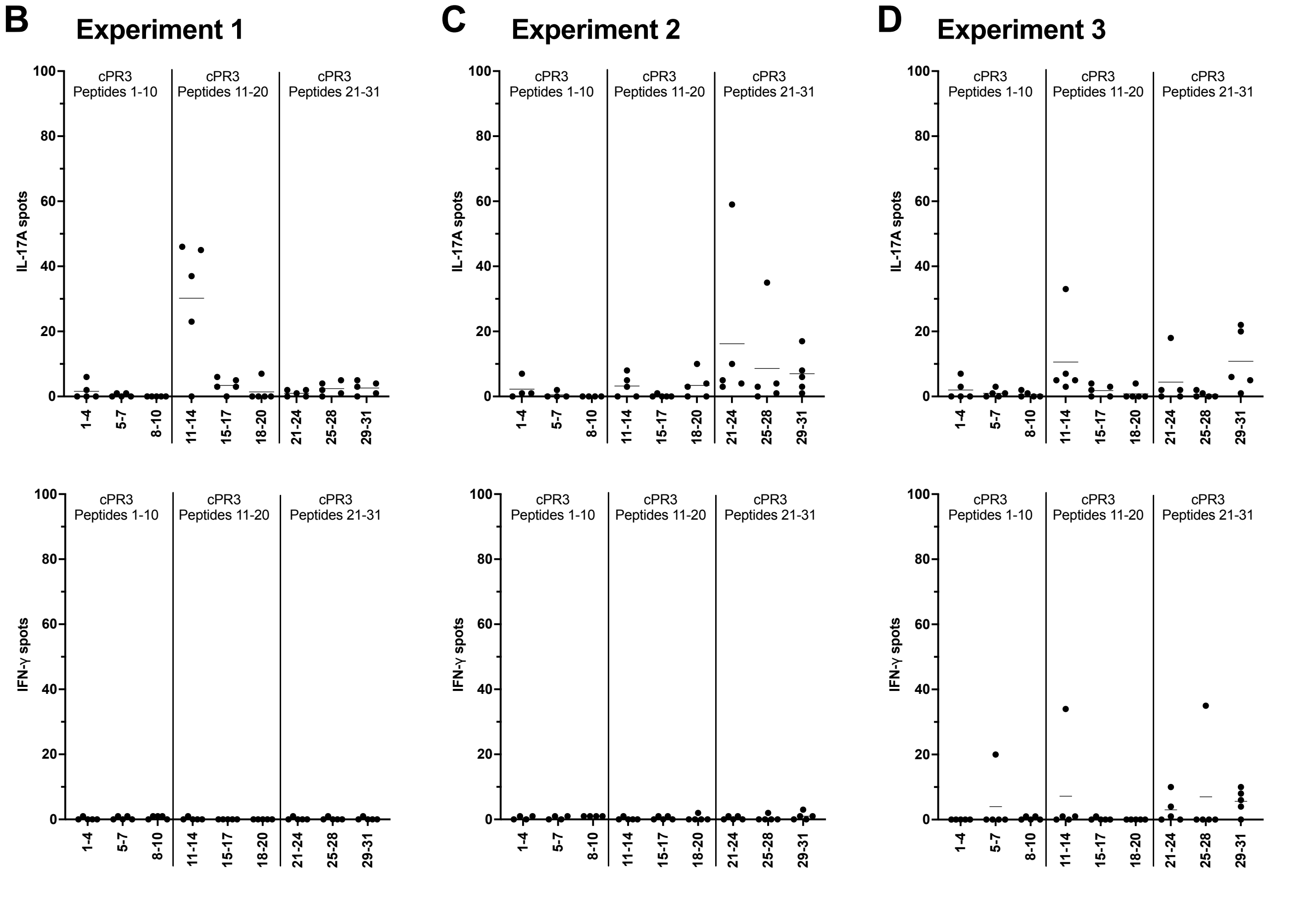


**Fig. S4. Lack of consistent reactivity to complementary PR3 (cPR3) peptides**

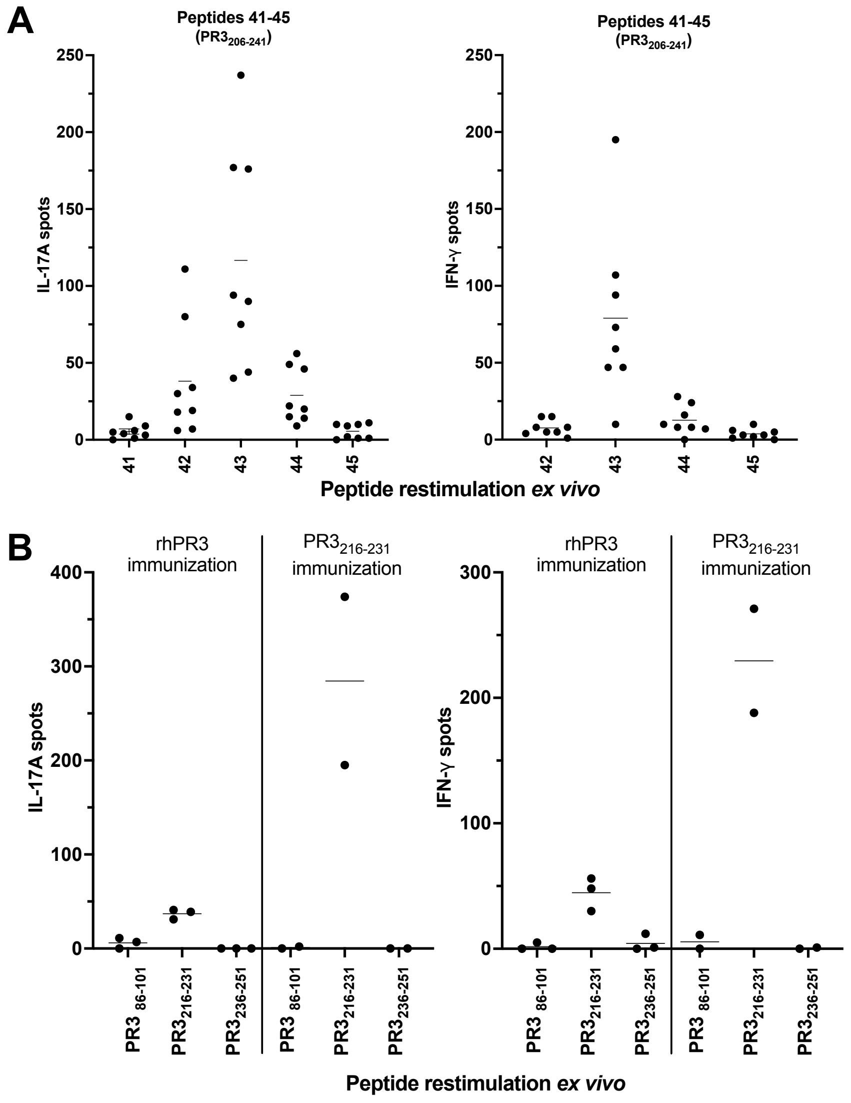


**Fig. S5. hPR3 peptide 43 ($PR3_{216-231}$) is confirmed as the immunodominant epitope, whereas peptide 47 ($PR3_{236-251}$) is non-immunogenic.**

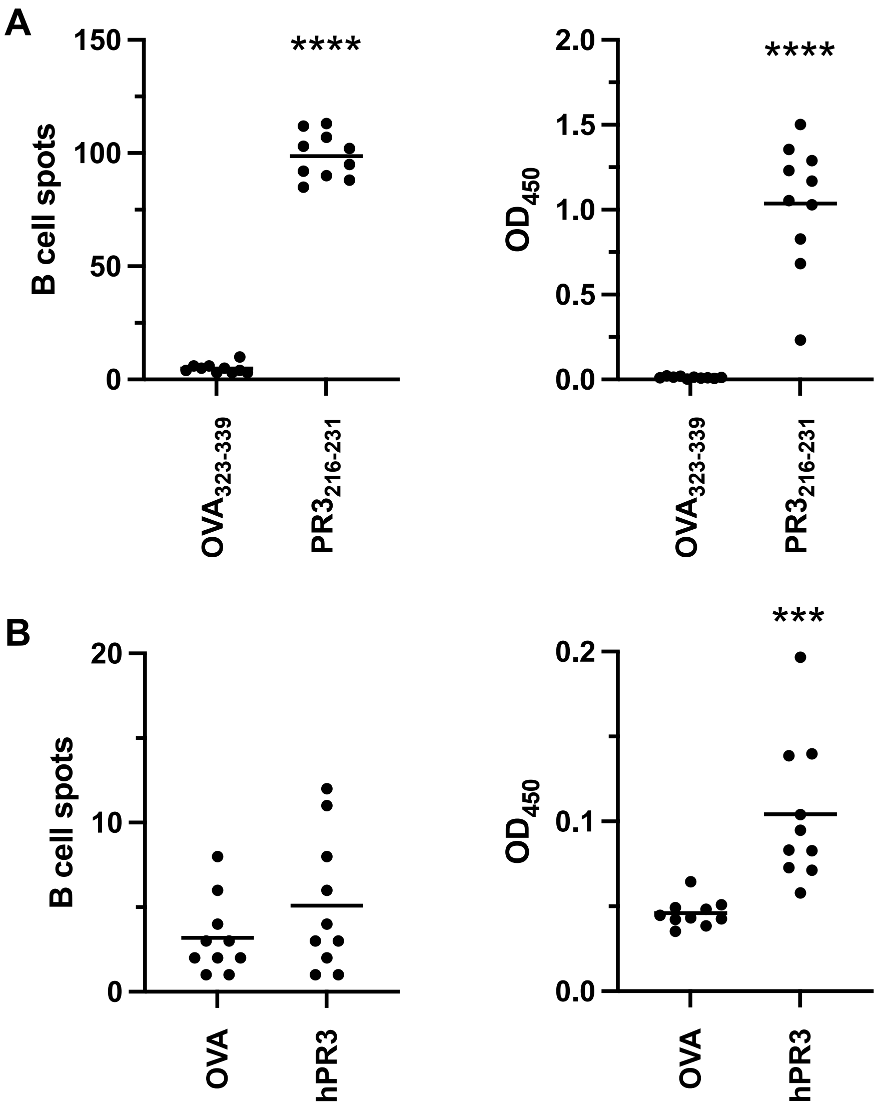


**Fig. S6. $PR3_{216\text{-}231}$ induces autoreactive B cell response.**

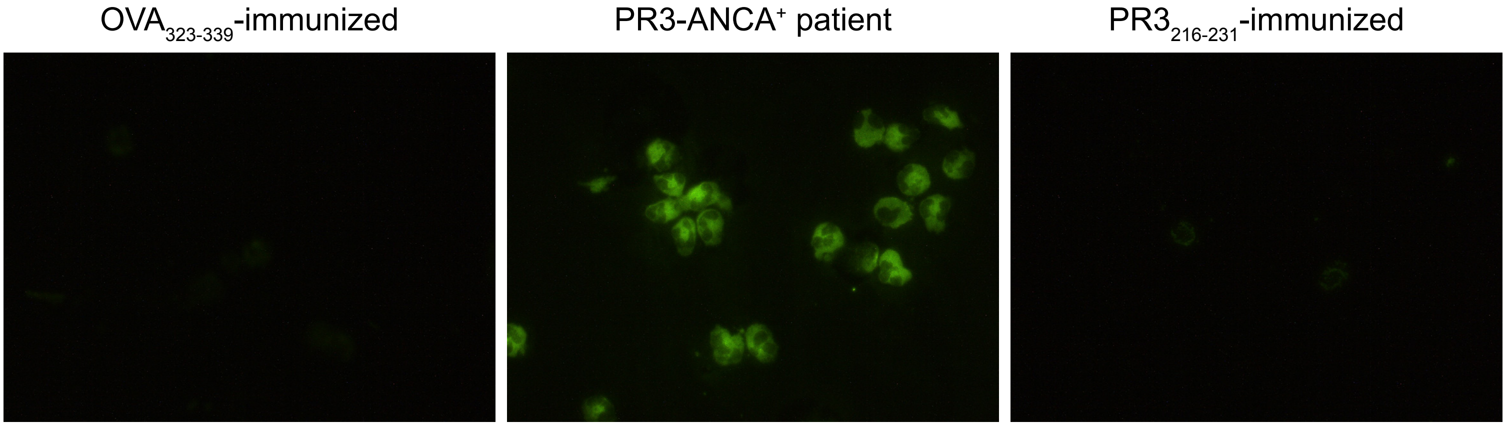


**Fig. S7. Antibodies from the sera of $PR3_{216\text{-}231}$ immunized hPR3.DR15$^{+}$ mice do not induce a cANCA pattern in human neutrophils.**

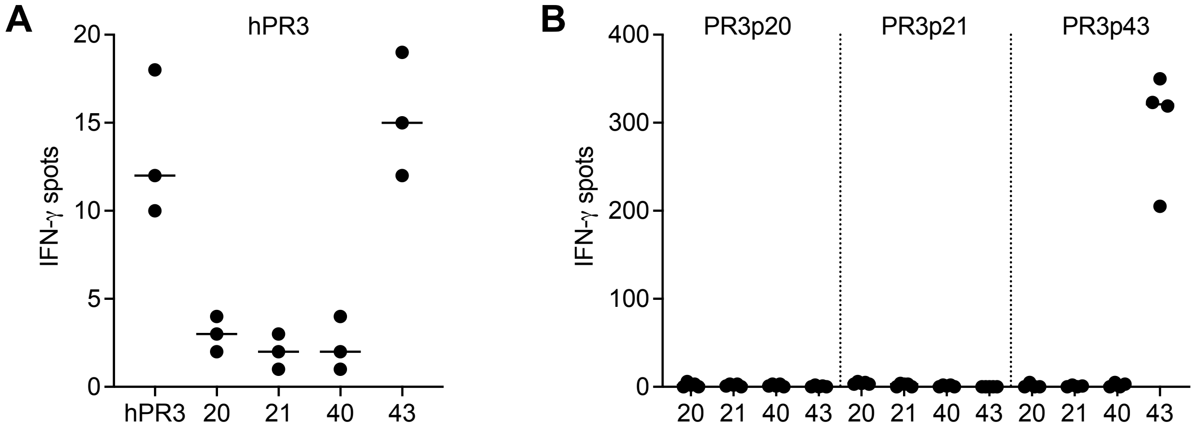


**Fig. S8. HLA-DR15-pressented peptide derived from the hPR3$_{103\text{-}119}$ region is non-immunogenic**

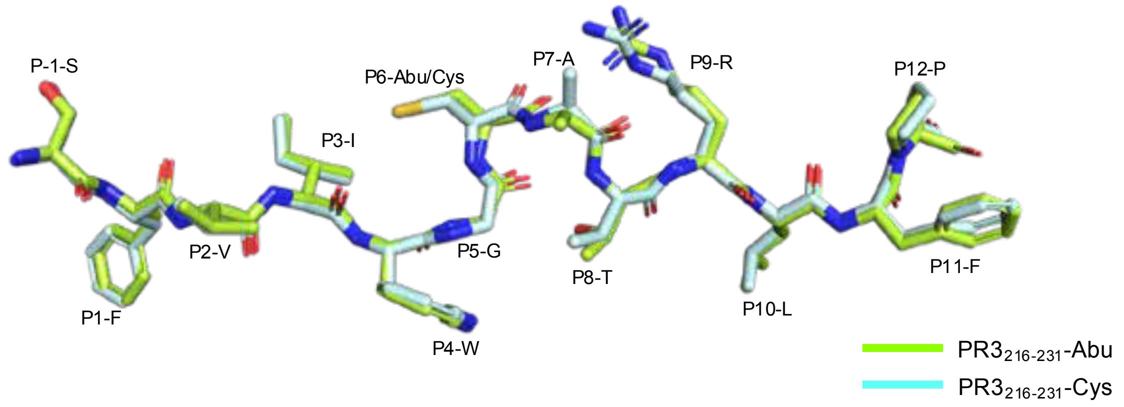


**Fig. S9. Superposition of HLA-DR15 presenting $PR3_{216\text{-}231}$Abu and $PR3_{216\text{-}231}$Cys.**

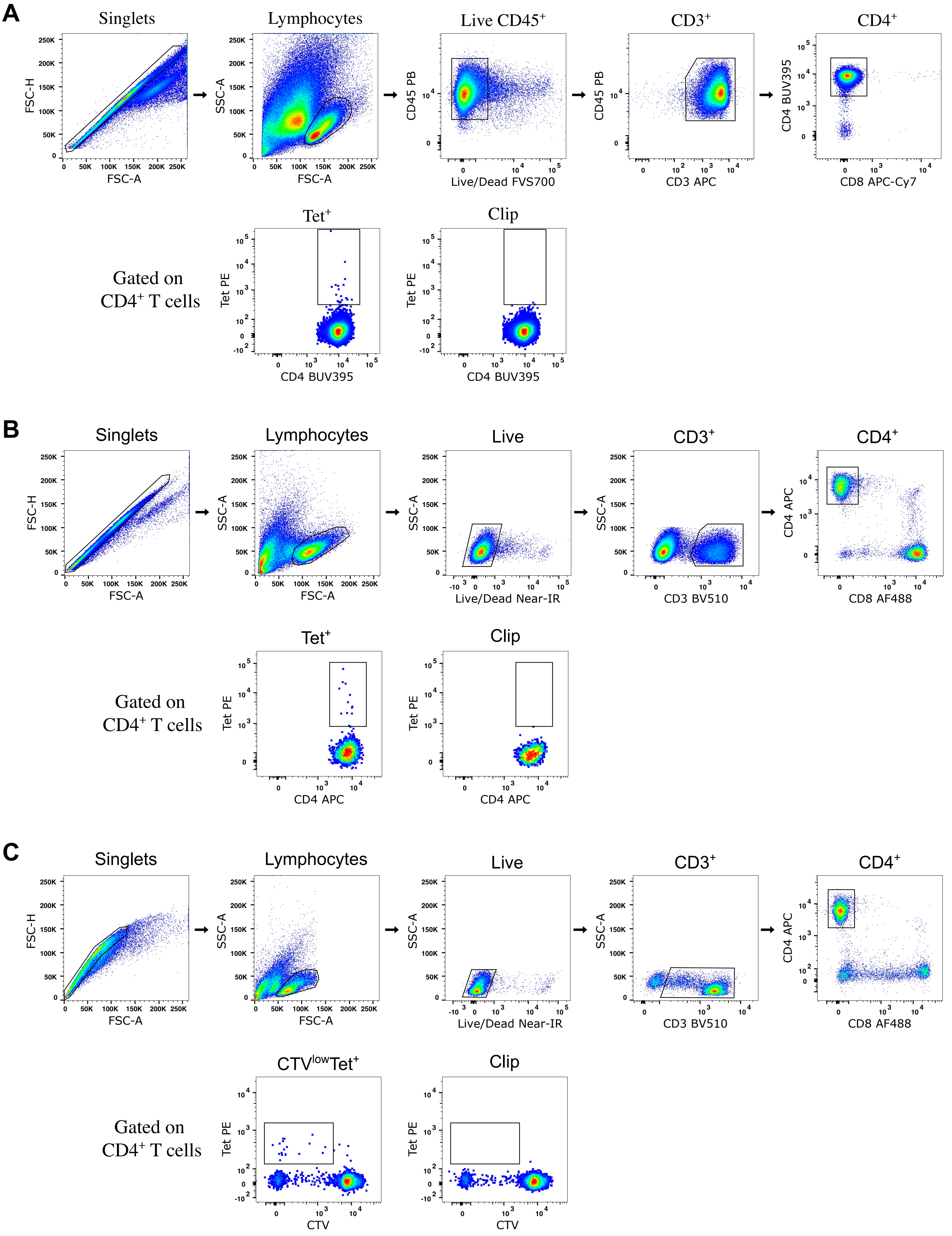


**Fig. S10. Flow cytometric gating strategies for single-cell sorting of DR15-PR3$_{216\text{-}231}$-tetramer$^+$ CD4$^+$ T cells.**

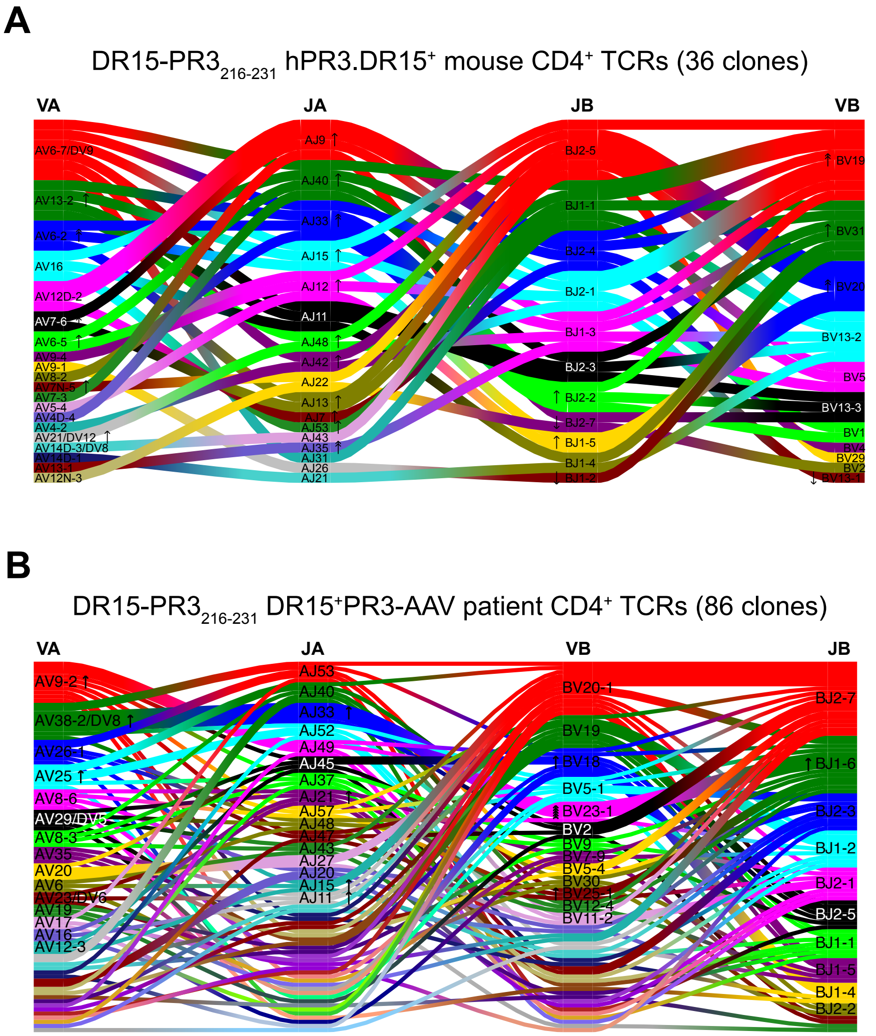


**Fig. S11. DR15-PR3$_{216\text{-}231}$-specific CD4$^{+}$ TCR repertoires in humanized hPR3.DR15$^{+}$ mice and DR15$^{+}$ PR3-AAV patients.**

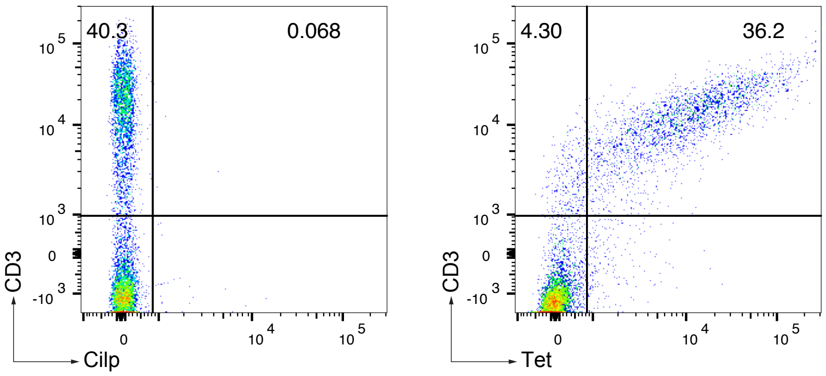


**Fig. S12. HLA-DR15-PR3$_{216\text{-}231}$ tetramer staining of cell line expressing TRAV16_TRBV19 TCR.**

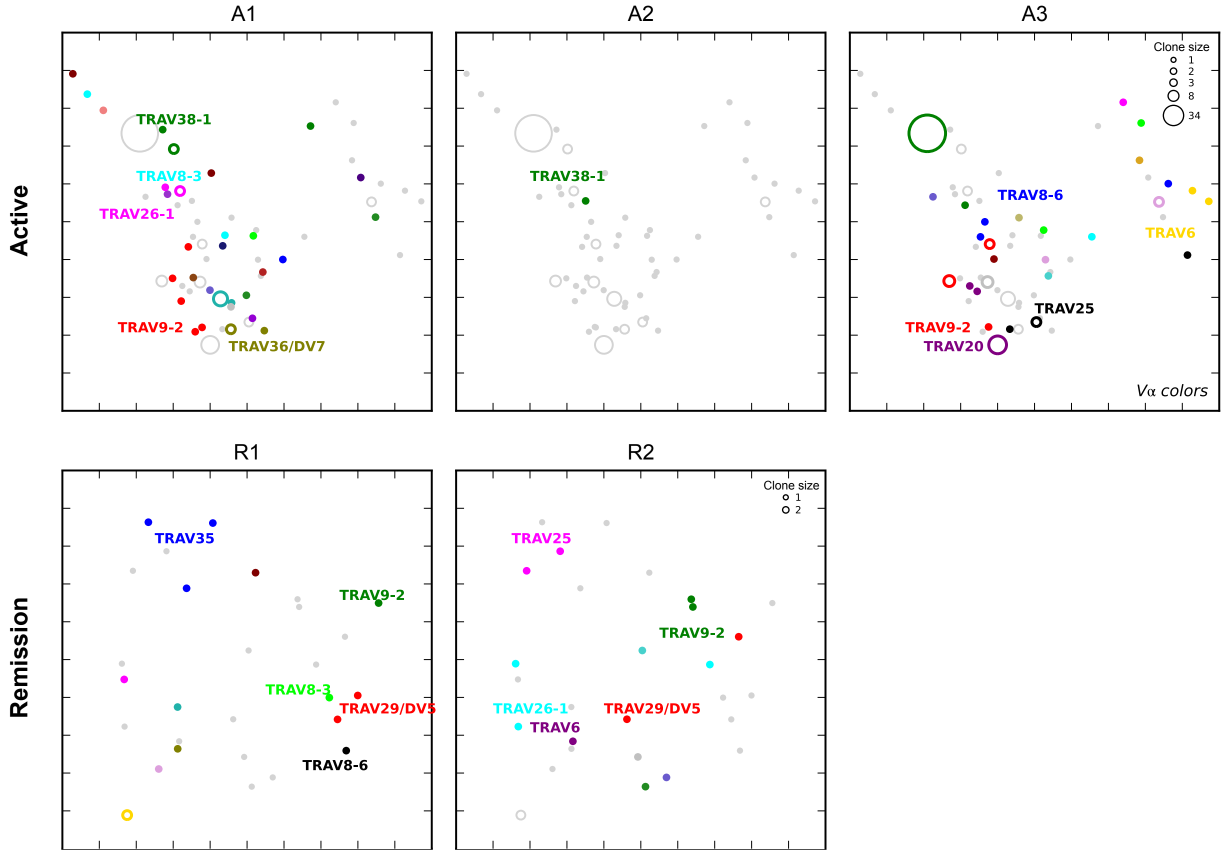


**Fig. S13. DR15-PR3$_{216\text{-}231}$-specific CD4$^{+}$ TCR clonotypes and their expansion in individual DR15$^{+}$ PR3-AAV patients.**

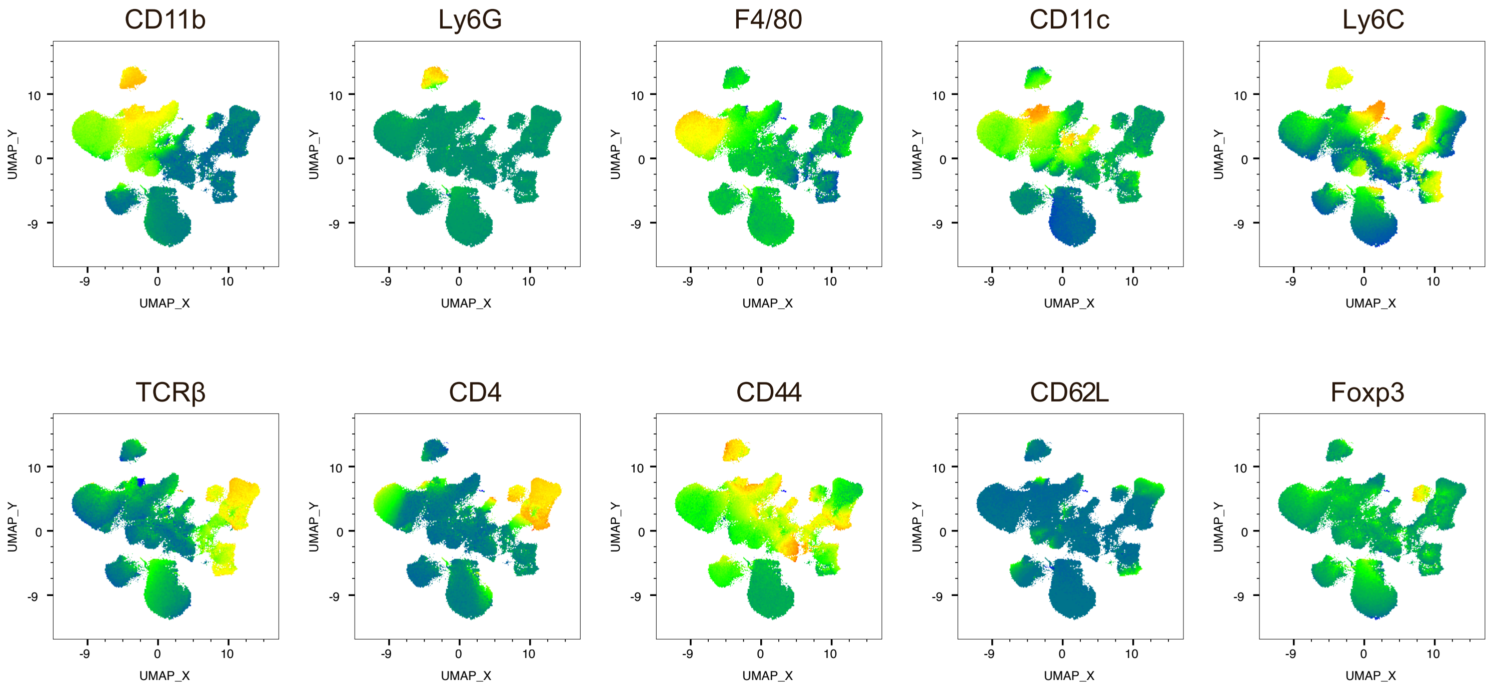


**Fig. S14. UMAP analysis of kidney leukocytes.**
