## Supplementary Tables 1-8 for "Unconventional presentation of an immunodominant HLA-DR15-restricted nephritogenic proteinase 3 epitope implicated in vasculitis"

**Table S1. Human PR3 (exons 2-5) peptides for CD4 T cell epitope screening**

| Peptide ID | Peptide sequence | Amino acid position |
|---|---|---|
| 4 | GAARAAEIVGGHEAQP | 21-36 |
| 5 | AEIVGGHEAQPHSRPY | 26-41 |
| 6 | GHEAQPHSRPYMASLQ | 31-46 |
| 7 | PHSRPYMASLQMRGNP | 36-51 |
| 8 | YMASLQMRGNPGSHFC | 41-56 |
| 9 | QMRGNPGSHFCGGTLI | 46-61 |
| 10 | PGSHFCGGTLIHPSFV | 51-66 |
| 11 | CGGTLIHPSFVLTAAH | 56-71 |
| 12 | IHPSFVLTAAHCLRDI | 61-76 |
| 13 | VLTAAHCLRDIPQRLV | 66-81 |
| 14 | HCLRDIPQRLVNVVLG | 71-86 |
| 15 | IPQRLVNVVLGAHNVR | 76-91 |
| 16 | VNVVLGAHNVRTQEPT | 81-96 |
| 17 | GAHNVRTQEPTQQHFS | 86-101 |
| 18 | RTQEPTQQHFSVAQVF | 91-106 |
| 19 | TQQHFSVAQVFLNNYD | 96-111 |
| 20 | SVAQVFLNNYDAENKL | 101-116 |
| 21 | FLNNYDAENKLNDVLL | 106-121 |
| 22 | DAENKLNDVLLIQLSS | 111-126 |
| 23 | LNDVLLIQLSSPANLS | 116-131 |
| 24 | LIQLSSPANLSASVAT | 121-136 |
| 25 | SPANLSASVATVQLPQ | 126-141 |
| 26 | SASVATVQLPQQDQPV | 131-146 |
| 27 | TVQLPQQDQPVPHGTQ | 136-151 |
| 28 | QQDQPVPHGTQCLAMG | 141-156 |
| 29 | VPHGTQCLAMGWGRVG | 146-161 |
| 30 | QCLAMGWGRVGAHDPP | 151-166 |
| 31 | GWGRVGAHDPPAQVLQ | 156-171 |
| 32 | GAHDPPAQVLQELNVT | 161-176 |
| 33 | PAQVLQELNVTVVTFF | 166-181 |
| 34 | QELNVTVVTFFCRPHN | 171-186 |
| 35 | TVVTFFCRPHNICTFV | 176-191 |
| 36 | FCRPHNICTFVPRRKA | 181-196 |
| 37 | NICTFVPRRKAGICFG | 186-201 |
| 38 | VPRRKAGICFGDSGGP | 191-206 |
| 39 | AGICFGDSGGPLICDG | 196-211 |
| 40 | GDSGGPLICDGIIQGI | 201-216 |
| 41 | PLICDGIIQGIDSFVI | 206-221 |
| 42 | GIIQGIDSFVIWGCAT | 211-226 |
| 43 | IDSFVIWGCATRLFPD | 216-231 |
| 44 | IWGCATRLFPDFFTRV | 221-236 |
| 45 | TRLFPDFFTRVALYVD | 226-241 |
| 46 | DFFTRVALYVDWIRST | 231-246 |
| 47 | VALYVDWIRSTLRRVE | 236-251 |
| 48 | DWIRSTLRRVEAKGRP | 241-256 |

**Table S2. Mouse and human PR3 (exon 1) peptides for T cell epitope screening**

| Peptide ID | Peptide sequence | Amino acid position |
|---|---|---|
| Mouse *Prtn3* exon 1 | | |
| M1 | MAGSYPSPKGIHPFLL | 1-16 |
| M2 | PSPKGIHPFLLLALVV | 5-21 |
| M3 | IHPFLLLALVVGGGAA | 10-26 |
| Human *PRTN3* exon 1 | | |
| H1 | MAHRPPSPALASVLLA | 1-16 |
| H2 | PSPALASVLLALLLSG | 5-21 |

**Table S3. Human complementary PR3 peptides for T cell epitope screening**

| Peptide ID | Peptide sequence | Amino acid position |
|---|---|---|
| Complementary Pool 1 | | |
| 1 | MWPAEEGDHGDIELLQ | 185-170 |
| 2 | EGDHGDIELLQDLGWG | 180-165 |
| 3 | DIELLQDLGWGVVGTH | 175-160 |
| 4 | QDLGWGVVGTHAAPAH | 170-155 |
| 5 | GVVGTHAAPAHGQALG | 165-150 |
| 6 | HAAPAHGQALGAVGHW | 160-145 |
| 7 | HGQALGAVGHWLVLLW | 155-140 |
| 8 | GAVGHWLVLLWQLDCG | 150-135 |
| 9 | WLVLLWQLDCGDGGTE | 145-130 |
| 10 | WQLDCGDGGTEVGWAA | 140-125 |
| Complementary Pool 2 | | |
| 11 | GDGGTEVGWAAQLDEE | 135-120 |
| 12 | EVGWAAQLDEENVVQF | 130-115 |
| 13 | AQLDEENVVQFVLRVV | 125-110 |
| 14 | ENVVQFVLRVVVVQKH | 120-105 |
| 15 | FVLRVVVVQKHLSHRE | 115-100 |
| 16 | VVVQKHLSHREVLLGG | 110-95 |
| 17 | HLSHREVLLGGLLRPH | 105-90 |
| 18 | EVLLGGLLRPHVVGSE | 100-85 |
| 19 | GLLRPHVVGSEHHVHQ | 95-80 |
| 20 | HVVGSEHHVHQALGYV | 90-75 |
| Complementary Pool 3 | | |
| 21 | EHHVHQALGYVPQAVR | 85-70 |
| 22 | QALGYVPQAVRGRQHE | 80-65 |
| 23 | VPQAVRGRQHEAGVDQ | 75-60 |
| 24 | RGRQHEAGVDQGASAE | 70-55 |
| 25 | EAGVDQGASAEVAARV | 65-50 |
| 26 | QGASAEVAARVPPHLQ | 60-45 |
| 27 | EVAARVPPHLQGGHVG | 55-40 |
| 28 | VPPHLQGGHVGPGVWL | 50-35 |
| 29 | QGGHVGPGVWLRLVPA | 45-30 |
| 30 | GPGVWLRLVPAHDLRS | 40-25 |
| 31 | LRLVPAHDLRSSGSTA | 35-20 |

**Table S4. Data collection and refinement statistics of HLA-DR15-PR3 binary structures**

| | DR15- $PR3_{216-231}$-Abu | DR15- $PR3_{216-231}$-Cys |
|---|---|---|
| **Data collection** | | |
| Space group | P6122 | P6122 |
| Cell dimensions | | |
| a, b, c (Å) | 147.40, 147.40, 139.27 | 146.95, 146.95, 139.49 |
| $\alpha$, $\beta$, $\gamma$ (°) | 90, 90, 120 | 90, 90, 120 |
| Resolution (Å)[a] | 47.05-2.88 (3.03-2.88) | 47.01-3.00 (3.18-3.00) |
| $R_{sym}$ or $R_{merge}$ [a] | 0.10 (1.09) | 0.11 (1.20) |
| I /$\sigma$ (I)[a] | 28.6 (4.2) | 22.4 (3.3) |
| $CC_{1/2}$ [a] | 1.0 (0.95) | 1.0 (0.88) |
| Completeness (%)[a] | 99.9 (99.4) | 100.0 (100.0) |
| Redundancy [a] | 27.8 (28.6) | 19.9 (20.5) |
| **Refinement** | | |
| Resolution (Å) | 47.05-2.88 | 47.01-3.00 |
| No. reflections | 20790 | 18332 |
| $R_{work}$ / $R_{free}$ | 0.247/0.205 | 0.241/0.209 |
| No. atoms | 3276 | 3263 |
| Protein | 3168 | 3140 |
| Ligand/ion | 72 | 102 |
| Water | 36 | 21 |
| B-factors (Å$^2$) | 76.5 | 80.4 |
| Protein | 75.7 | 79.1 |
| Ligand/ion | 116.4 | 123.4 |
| Water | 69.1 | 77.6 |
| Ramachandran | | |
| Outlier (%) | 0.00 | 0.00 |
| Favored (%) | 95.54 | 96.07 |
| R.m.s. deviations | | |
| Bond lengths (Å) | 0.006 | 0.004 |
| Bond angles (°) | 0.775 | 0.603 |

[a] Values in parentheses refer to the highest resolution shell

#### Table S5. DR15-PR3$_{216-231}$-specific TCRαβ repertoire from PR3$_{216-231}$-immunized hPR3.DR15$^{+}$ mice

| TRAV | TRAJ | TRBV | TRBJ | TRBD | CDR3α | CDR3β | No. Clone | Frequency |
|---|---|---|---|---|---|---|---|---|
| 6-7/DV9 | 42 | 31 | 1-1 | 1 | SAGGSNAKL | SLGAGEV | 3 | 8 |
| 16 | 53 | 19 | 1-1 | 1 | REGGGSNYKL | SIWGEV | 2 | 5 |
| 13-2 | 15 | 20 | 2-5 | 2 | YQGGRAL | RTGGQDTQ | 2 | 5 |
| 4-2 | 42 | 13-1 | 2-1 | 2 | GGSNAKL | RDNYAEQ | 1 | 3 |
| 4-4/DV10 | 40 | 13-2 | 1-3 | 1 | NTGNYKY | GQGGGNTL | 1 | 3 |
| 5D-4 | 12 | 20 | 1-5 | 1 | HQTGGYKV | RERVNQAP | 1 | 3 |
| 6-3 | 15 | 13-2 | 1-5 | 1 | REPQGGRAL | GRGYNNQAP | 1 | 3 |
| 6-3 | 13 | 19 | 2-5 | 2 | RNSGTYQ | SIAGSQDTQ | 1 | 3 |
| 6-3 | 33 | 19 | 2-2 | 1 | RDRSNYQL | SIPGQL | 1 | 3 |
| 6-5 | 12 | 1 | 2-1 | 1 | SDGYKV | SADPAGPNYAEQ | 1 | 3 |
| 6-5 | 40 | 5 | 1-1 | 1 | SDAGNYKY | SLRGRFTEV | 1 | 3 |
| 6-6 | 40 | 4 | 2-4 | 2 | GDPGNYKY | SSGTGGNTL | 1 | 3 |
| 6-7/DV9 | 11 | 13-3 | 2-3 | 2 | DSGYNKL | RGLGGRGAETL | 1 | 3 |
| 6-7/DV9 | 15 | 5 | 1-3 | 1 | GPYQGGRAL | SHPPGQGAGTL | 1 | 3 |
| 6-7/DV9 | 31 | 13-2 | 1-1 | 1 | GDSNNRI | GDAGQGPANTEV | 1 | 3 |
| 6D-7 | 33 | 13-3 | 2-2 | 1 | GVDSNYQL | SAPGQGYTGQL | 1 | 3 |
| 7D-3 | 9 | 19 | 2-1 | 2 | SLNMGYKL | GQNYAEQ | 1 | 3 |
| 7D-5 | 22 | 13-3 | 2-7 | 2 | SLASSGSWQL | SDWGGDEQ | 1 | 3 |
| 7D-6 | 9 | 5 | 2-4 | 2 | NMGYKL | SQEGNTL | 1 | 3 |
| 7D-6 | 13 | 19 | 1-1 | 1 | SNSGTYQ | SIGFDTEV | 1 | 3 |
| 8-2 | 33 | 19 | 2-1 | 1 | DDSNYQL | TFYAEQ | 1 | 3 |
| 9-1 | 26 | 31 | 1-2 | 1 | SGNYAQGL | SLGNSDY | 1 | 3 |
| 9N-4 | 12 | 20 | 2-5 | 1 | SMVTGGYKV | RDRTGVNQDTQ | 1 | 3 |
| 12D-2 | 9 | 2 | 2-7 | 1 | SGNMGYKL | SQERVEQ | 1 | 3 |
| 12D-2 | 7 | 19 | 2-4 | 2 | KDYSNNRL | SPYWGQNTL | 1 | 3 |
| 12D-2 | 9 | 31 | 2-4 | 1 | SGMGYKL | SLGQGNTL | 1 | 3 |
| 13-1 | 35 | 19 | 1-3 | 1 | ATGFASAL | TGTGEGNTL | 1 | 3 |
| 13-2 | 11 | 13-2 | 2-3 | 2 | DRSGYNKL | GDGGTSAETL | 1 | 3 |
| 13-2 | 33 | 1 | 2-2 | 1 | DRSNYQL | SAQNTGQL | 1 | 3 |
| 13-2 | 48 | 31 | 1-3 | 1 | DHYGNEKI | RQGFGNTL | 1 | 3 |
| 14-3 | 43 | 29 | 2-5 | 1 | SVDNNAP | SSPPQDTQ | 1 | 3 |
| 14D-1 | 21 | 31 | 1-4 | 1 | SRGNYNVL | SLGTGEVERL | 1 | 3 |
| 14N-1 | 4 | 13-1 | 1-4 | 1 | SGLGGFNKL | SDGGFSNERL | 1 | 3 |
| 16D/DV11 | 40 | 31 | 2-3 | 2 | INTGNYKY | SWGGAETL | 1 | 3 |
| 16D/DV11 | 11 | 13-2 | 1-4 | 1 | KGSGYNKL | GTGTGVSNERL | 1 | 3 |
| 21 | 48 | 13-2 | 2-5 | 2 | TNYGNEKI | GDAGINQDTQ | 1 | 3 |

#### Table S6. DR15$^+$ PR3-AAV patient details

| | A1 | A2 | A3 | R1 | R2 |
|---|---|---|---|---|---|
| **Phenotype** | GPA | GPA | GPA | GPA | GPA |
| **Time from diagnosis to sampling (years)** | 7, 7.3 | At diagnosis | At diagnosis | 22 | 2.5, 2.7 |
| **Age at sampling (years)** | 66, 67 | 70 | 56 | 61 | 69, 70 |
| **PR3-ANCA (<20 IU/mL)** | >160 | >200 | >200 | 91 | 113 |
| **Organ/tissue involvement (any time)** | ENT, lung, kidney, PNS | ENT, kidney | Kidney, musculoskeletal | ENT, lung, musculoskeletal | ENT, lung, musculoskeletal |
| **Activity at sampling** | Relapse: ENT, kidney | Initial: ENT, kidney | Initial: kidney, musculoskeletal | Inactive: 5 years | Inactive: 12 months |
| **Treatment at sampling** | Methylprednisolone i.v. 1 dose | Methylprednisolone i.v. 3 doses | Methotrexate | Methotrexate | Methotrexate |

ENT, ear, nose and throat; GPA, granulomatosis with polyangiitis, PNS, peripheral nervous system

### Table S7. DR15-PR3$_{216\text{-}231}$-specific TCRαβ repertoire from DR15⁺ PR3-AAV patients

| TRAV | TRAJ | TRBV | TRBJ | TRBD | CDR3α | CDR3β | No. Clone | Frequency | Sample ID |
|---|---|---|---|---|---|---|---|---|---|
| 38-2/DV8 | 33 | 23-1 | 1-6 | 1 | RFMDSNYQL | SNRGGRATGHNSPL | 36 | 25 | R1(2), A3(34) |
| 20 | 27 | 19 | 1-2 | 1 | QAGTNAGKS | SWLSYGY | 9 | 6 | R1(1), A3(8) |
| 12-2 | 20 | 5-5 | 2-3 | 2 | SWYDYKL | SSGLAADTQ | 5 | 3 | A1 |
| 9-2 | 45 | 15 | 1-4 | 1 | KGGGADGL | SRDRTSNEKL | 3 | 2 | A3 |
| 12-1 | 37 | 11-2 | 2-7 | 2 | ARGSSNTGKL | SLDPQGGQ | 3 | 2 | A3 |
| 26-1 | 57 | 5-1 | 2-3 | 1 | RFVTQGGSEKL | SFTGGSVTQ | 2 | 1 | A1 |
| 1-2 | 12 | 6-4 | 2-3 | 2 | MDSSYKL | SDSSGGADTQ | 1 | 1 | R2 |
| 3 | 8 | 20-1 | 1-6 | 1 | FQKL | RFAARDNSPL | 1 | 1 | A1 |
| 4 | 5 | 19 | 1-1 | 1 | GLLDTGRRAL | KGKELNTEA | 1 | 1 | A3 |
| 5 | 9 | 7-6 | 2-2 | 1 | TSNTGGFKT | SLSHNVPGEL | 1 | 1 | A1 |
| 6 | 15 | 20-1 | 2-7 | 2 | ENQAGTAL | RAGWLAYEQ | 1 | 1 | A3 |
| 6 | 43 | 20-1 | 2-7 | 2 | VNNNDM | RTSGRIYEQ | 1 | 1 | A3 |
| 6 | 6 | 7-9 | 1-2 | 1 | DFSGGSYIP | SLGTPPNYGY | 1 | 1 | R2 |
| 8-2 | 3 | 10-3 | 1-3 | 1 | EYSSASKI | RDTFSGNTI | 1 | 1 | A3 |
| 8-3 | 45 | 18 | 1-4 | 1 | GPNSDSGGGADGL | SPDGLTNEKL | 1 | 1 | A1 |
| 8-3 | 40 | 2 | 2-7 | 2 | GARGTYKY | RDPAYEQ | 1 | 1 | R1 |
| 8-3 | 37 | 29-1 | 2-3 | 1 | SNTGKL | EGHGDTQ | 1 | 1 | A3 |
| 8-3 | 43 | 7-3 | 2-1 | 2 | VPYNNNDM | SLNKGPNNEQ | 1 | 1 | A1 |
| 8-6 | 20 | 10-3 | 1-1 | 1 | ISSNDYKL | RGTGRNTEA | 1 | 1 | A1 |
| 8-6 | 22 | 15 | 1-1 | 1 | TVSGAARQL | SRDRTNTEA | 1 | 1 | A3 |
| 8-6 | 57 | 19 | 1-2 | 1 | SERRGSEKL | SMGMDRGPANYGY | 1 | 1 | A3 |
| 8-6 | 24 | 20-1 | 2-7 | 1 | MTTDSWGKL | RARDRGYEQ | 1 | 1 | A3 |
| 8-6 | 21 | 27 | 1-1 | 1 | SKYNFNKF | WAFEDRVNTEA | 1 | 1 | R1 |
| 9-2 | 42 | 19 | 1-5 | 1 | GGSQGNL | SSTGSGQPQ | 1 | 1 | R2 |
| 9-2 | 47 | 2 | 2-7 | 1 | SNYSHGNKL | SVRYEQ | 1 | 1 | A3 |
| 9-2 | 57 | 25-1 | 2-1 | 2 | SRQGGSEKL | SELYEQ | 1 | 1 | A1 |
| 9-2 | 49 | 29-1 | 2-3 | 2 | SSNTGNQF | GGVSSDTQ | 1 | 1 | R1 |
| 9-2 | 49 | 3-1 | 2-1 | 2 | SDPRTNTGNQF | SHPEGEQ | 1 | 1 | A1 |
| 9-2 | 24 | 30 | 1-5 | 1 | RIMTTDSWGKL | TPPGGHQPQ | 2 | 1 | A3 |
| 9-2 | 54 | 4-1 | 1-4 | 1 | STQGAQKL | SQGSLGNEKL | 1 | 1 | A1 |
| 9-2 | 49 | 5-1 | 2-5 | 1 | SDPSTGNQF | SLGMETQ | 1 | 1 | R2 |
| 9-2 | 7 | 5-1 | 1-6 | 1 | SDRDGNNRL | RPQGTYSPL | 1 | 1 | A1 |
| 9-2 | 37 | 9 | 2-5 | 1 | GSSNTGKL | SVGWTGQETQ | 1 | 1 | A1 |
| 12-1 | 36 | 19 | 2-7 | 2 | NIPGANNL | SIERIYEQ | 1 | 1 | A1 |
| 12-2 | 20 | 25-1 | 1-6 | 1 | SWYDYKL | SESRGRDSPL | 1 | 1 | A1 |
| 12-3 | 45 | 18 | 2-5 | 2 | SESGGGADGL | SPRGETQ | 1 | 1 | A3 |
| 12-3 | 15 | 5-5 | 1-1 | 1 | FNQAGTAL | SNAALRNTEA | 1 | 1 | R2 |
| 12-3 | 21 | 7-8 | 1-2 | 1 | DVYNFNKF | SDGQGKAGY | 1 | 1 | A1 |
| 13-1 | 13 | 20-1 | 2-5 | 2 | SSGGYQKV | RDRRSPLETQ | 2 | 1 | A3 |
| 13-1 | 52 | 30 | 2-6 | 2 | GGTSYGKL | SGAGANVL | 1 | 1 | A3 |
| 13-2 | 9 | 20-1 | 1-5 | 1 | NPKTGGFKT | RQDPSNQPQ | 1 | 1 | A3 |
| 14/DV4 | 47 | 5-1 | 2-3 | 2 | RVIMEYGNKL | SLVGAFTDTQ | 1 | 1 | R1 |
| 16 | 11 | 19 | 2-5 | 1 | TGYSTL | SPARGLATQ | 1 | 1 | A1 |
| 16 | 29 | 20-1 | 2-7 | 2 | LSGNTPL | PRLAGGSYEQ | 1 | 1 | A1 |
| 16 | 21 | 7-9 | 1-6 | 1 | DNFNKF | SLGLGSYNSPL | 1 | 1 | R2 |
| 17 | 53 | 18 | 1-4 | 1 | GRVNSGGSNYKL | SPDGLTNEKL | 1 | 1 | A1 |
| 17 | 16 | 7-9 | 2-2 | 1 | AGEFSDGQKL | SPNRGHANTGEL | 1 | 1 | R1 |
| 19 | 28 | 18 | 1-4 | 1 | SDKAYSGAGSYQL | SPDGLTNEKL | 1 | 1 | A1 |
| 19 | 31 | 2 | 2-6 | 2 | SGYNNARL | SGRALGANVL | 1 | 1 | R2 |
| 19 | 52 | 6-5 | 2-7 | 1 | ALPAGGTSYGKL | RRDRGSYEQ | 1 | 1 | A1 |
| 20 | 45 | 5-4 | 2-7 | 2 | QAGGGADGL | SAGLAKYEQ | 1 | 1 | A3 |
| 20 | 27 | 7-6 | 1-6 | 1 | QAGTNAGKS | SPGRAYNSPL | 1 | 1 | A3 |
| 21 | 5 | 30 | 2-3 | 1 | STGRRAL | SAGTGETDTQ | 1 | 1 | A1 |
| 21 | 28 | 9 | 2-1 | 2 | GGYSGAGSYQL | SVASGMGNEQ | 1 | 1 | R1 |
| 23/DV6 | 17 | 12-5 | 2-3 | 2 | SIRAAGNKL | GLGGGPDTQ | 1 | 1 | A1 |
| 23/DV6 | 11 | 20-1 | 1-6 | 1 | SVSEGSGYSTL | RVSLDRGSPL | 1 | 1 | A3 |
| 23/DV6 | 48 | 27 | 1-2 | 1 | SGDFGNEKL | RASRGTYYYGY | 1 | 1 | A3 |
| 25 | 40 | 11-2 | 2-1 | 1 | GTGTYKY | SLLGQSNEQ | 2 | 1 | A3 |
| 25 | 40 | 20-1 | 2-7 | 2 | GTGTYKY | RTSGRIYEQ | 1 | 1 | A3 |
| 25 | 52 | 4-1 | 1-2 | 1 | SRSGTSYGKL | SPLGRNGY | 1 | 1 | R2 |
| 25 | 32 | 5-1 | 2-2 | 2 | PPYGGATNKL | SIGTTNTGEL | 1 | 1 | R1 |
| 25 | 37 | 6-2 | 2-1 | 2 | KSSNTGKL | SYVGERG | 1 | 1 | A3 |
| 25 | 52 | 6-4 | 1-5 | 1 | KVGTSYGKL | SVEYNQPQ | 1 | 1 | R2 |
| 26-1 | 53 | 12-3 or 12-4 | 1-5 | 1 | RIPLSGGSRGNM | ASTGPNQPQ | 1 | 1 | R2 |
| 26-1 | 26 | 20-1 | 2-7 | 2 | NYGQNF | RRRAFRTEYEQ | 1 | 1 | R2 |
| 26-1 | 15 | 20-1 | 1-2 | 1 | SPLGQAGTAL | RGDREGRGGY | 1 | 1 | A3 |
| 26-1 | 48 | 23-1 | 1-2 | 1 | RGSSNFGNEKL | SQSGRNYGY | 1 | 1 | R2 |
| 26-1 | 53 | 5-4 | 2-7 | 2 | RDRGGSNYKL | SPERREQ | 1 | 1 | A1 |
| 27 | 17 | 6-6 | 2-7 | 2 | ARAAGNKL | SFAGYEQ | 1 | 1 | A1 |
| 29/DV5 | 43 | 11-2 | 2-1 | 2 | SADNNDM | SFSETYNEQ | 1 | 1 | R1 |
| 29/DV5 | 21 | 12-3 or 12-4 | 2-7 | 2 | SGNFNKF | TSKASYEQ | 1 | 1 | R1 |
| 29/DV5 | 39 | 18 | 1-6 | 1 | ALNNAGNML | SLSRGLNDSPL | 1 | 1 | A1 |
| 29/DV5 | 36 | 19 | 2-5 | 2 | SARSGANNL | RHRGTTQETQ | 1 | 1 | R2 |
| 29/DV5 | 48 | 9 | 1-6 | 1 | ILNFGNEKL | SARDNSPL | 1 | 1 | R2 |
| 35 | 35 | 18 | 1-1 | 1 | QTGFGNVL | SYRVANTEA | 1 | 1 | A1 |
| 35 | 54 | 20-1 | 1-5 | 1 | QLRGGAQKL | LTGNQPQ | 1 | 1 | R1 |
| 35 | 15 | 28 | 1-3 | 1 | PNQAGTAL | SPHPLGLTGRNTI | 1 | 1 | R1 |
| 35 | 42 | 5-4 | 2-3 | 2 | ARPLNYGGSQGNL | SDRDTQ | 1 | 1 | R1 |
| 36/DV7 | 40 | 25-1 | 1-6 | 1 | PRGGTYKY | SESRGRDSPL | 2 | 1 | A1 |
| 36/DV7 | 40 | 6-1 | 1-1 | 1 | PRGGTYKY | RARGGTEA | 1 | 1 | A1 |
| 38-1 | 53 | 7-2 | 2-5 | 2 | REGGGSNYKL | SLDLAGGRETQ | 1 | 1 | A1 |
| 38-2/DV8 | 47 | 12-3 or 12-4 | 1-6 | 1 | RLLYGNKL | SLEGGGSPL | 1 | 1 | A2 |
| 38-2/DV8 | 33 | 2 | 2-7 | 1 | RFMDSNYQL | SVRYEQ | 1 | 1 | A3 |
| 38-2/DV8 | 53 | 20-1 | 2-1 | 1 | RKHGGRTDKS | GQTLYNEQ | 1 | 1 | A1 |
| 38-2/DV8 | 52 | 5-1 | 2-7 | 2 | RPRAGGTSYGKL | SSGLVGYEQ | 2 | 1 | A1 |
| 41 | 49 | 18 | 2-7 | 2 | LGNQF | SPQGGSLREQ | 1 | 1 | A3 |

#### Table S8. Antibodies and reagents used in flow cytometry

| Mouse Antibody (Clone) | Source | Catalogue number |
|---|---|---|
| Purified Rat Anti-Mouse CD16/CD32 (2.4G2) | BD Bioscience | 553142 |
| PerCP/Cyanine5.5 anti-mouse TCR β chain (H57-597) | Biolegend | 109228 |
| PE-Cy™7 Hamster Anti-Mouse CD69 (H1.2F3) | BD Bioscience | 552879 |
| Pacific Blue™ anti-mouse CD45 (30-F11) | Biolegend | 103126 |
| FITC Rat Anti-Mouse Ly-6C (AL-21) | BD Bioscience | 553104 |
| eFluor™ 450 FOXP3 Monoclonal Antibody (FJK-16s) | Thermo Fisher Scientific | 48-5773-82 |
| BV480 Rat Anti-Mouse CD62L (MEL-14 ) | BD Bioscience | 746726 |
| BUV805 Rat Anti-CD11b (M1/70) | BD Bioscience | 741934 |
| BUV737 Hamster Anti-Mouse CD11c (N418) | BD Bioscience | 569236 |
| BUV661 Rat Anti-Mouse CD4 (GK1.5) | BD Bioscience | 612974 |
| BUV496 Rat Anti-Mouse CD44 (IM7) | BD Bioscience | 741057 |
| BUV395 Rat Anti-Mouse CD8a (53-6.7) | BD Bioscience | 563786 |
| BUV395 Rat Anti-Mouse CD4 (GK1.5) | BD Bioscience | 563790 |
| Brilliant Violet 711™ anti-mouse F4/80 (BM8) | Biolegend | 123147 |
| Brilliant Violet 605™ anti-mouse Ly-6G (1A8) | Biolegend | 127639 |
| Brilliant Violet 510™ anti-mouse CD19 (6D5) | Biolegend | 115546 |
| APC-Cy™7 Rat Anti-Mouse CD8a (53-6.7) | BD Bioscience | 557654 |
| APC-Cy™7 Mouse Anti-Mouse CD45.2 (104) | BD Bioscience | 560694 |
| APC CD3e Monoclonal Antibody (145-2C11) | Thermo Fisher Scientific | 17-0031-82 |

| Human Antibody (Clone) | Source | Catalogue number |
|---|---|---|
| Brilliant Violet 510™ anti-human CD3 (OKT3) | Biolegend | 317332 |
| APC Mouse Anti-Human CD4 (SK3) | BD Bioscience | 566915 |
| Alexa Fluor® 488 anti-human CD8a (HIT8a) | Biolegend | 300916 |

| Reagent | Source | Catalogue number |
|---|---|---|
| Fixable Viability Stain 700 | BD Bioscience | 564997 |
| LIVE/DEAD™ Fixable Near-IR Dead Cell Stain Kit | Thermo Fisher Scientific | L10119 |
| CellTrace™ Violet Cell Proliferation Kit, for flow cytometry | Thermo Fisher Scientific | C34557 |
| eBioscience™ Foxp3 / Transcription Factor Staining Buffer Set | Thermo Fisher Scientific | 00-5523-00 |
